# A reduced Gompertz model for predicting tumor age using a population approach

**DOI:** 10.1101/670869

**Authors:** C. Vaghi, A. Rodallec, R. Fanciullino, J. Ciccolini, J. Mochel, M. Mastri, C. Poignard, J. ML Ebos, S. Benzekry

## Abstract

Tumor growth curves are classically modeled by ordinary differential equations. In analyzing the Gompertz model several studies have reported a striking correlation between the two parameters of the model.

We analyzed tumor growth kinetics within the statistical framework of nonlinear mixed-effects (population approach). This allowed for the simultaneous modeling of tumor dynamics and interanimal variability. Experimental data comprised three animal models of breast and lung cancers, with 843 measurements in 94 animals. Candidate models of tumor growth included the Exponential, Logistic and Gompertz. The Exponential and – more notably – Logistic models failed to describe the experimental data whereas the Gompertz model generated very good fits. The population-level correlation between the Gompertz parameters was further confirmed in our analysis (R^2^ > 0.96 in all groups). Combining this structural correlation with rigorous population parameter estimation, we propose a novel reduced Gompertz function consisting of a single individual parameter. Leveraging the population approach using bayesian inference, we estimated the time of tumor initiation using three late measurement timepoints. The reduced Gompertz model was found to exhibit the best results, with drastic improvements when using bayesian inference as compared to likelihood maximization alone, for both accuracy and precision. Specifically, mean accuracy was 12.1% versus 74.1% and mean precision was 15.2 days versus 186 days, for the breast cancer cell line.

These results offer promising clinical perspectives for the personalized prediction of tumor age from limited data at diagnosis. In turn, such predictions could be helpful for assessing the extent of invisible metastasis at the time of diagnosis.

**Author summary:** Mathematical models for tumor growth kinetics have been widely used since several decades but mostly fitted to individual or average growth curves. Here we compared three classical models (Exponential, Logistic and Gompertz) using a population approach, which accounts for inter-animal variability. The Exponential and the Logistic models failed to fit the experimental data while the Gompertz model showed excellent descriptive power. Moreover, the strong correlation between the two parameters of the Gompertz equation motivated a simplification of the model, the reduced Gompertz model, with a single individual parameter and equal descriptive power. Combining the mixed-effects approach with Bayesian inference, we predicted the age of individual tumors with only few late measurements. Thanks to its simplicity, the reduced Gompertz model showed superior predictive power. Although our method remains to be extended to clinical data, these results are promising for the personalized estimation of the age of a tumor from limited measurements at diagnosis. Such predictions could contribute to the development of computational models for metastasis.

## 1 Introduction

In the era of personalized oncology, mathematical modelling is a valuable tool for quantitative description of physiopathological phenomena [1, 2]. It allows for a better understanding of biological processes and to generate useful individual clinical predictions, for instance for personalized dose adaptation in cancer therapeutic menagement [3]. Tumor growth kinetics have been studied since several decades both clinically [4] and experimentally [5]. One of the main findings of these early studies is that tumor growth is not entirely exponential, provided it is observed over a long enough timeframe (100 to 1000 folds of increase) [6]. The specific growth rate slows down and this deceleration can be particularly well captured by the Gompertz model [7, 6, 8]. The analytical expression of this model reads as follows:

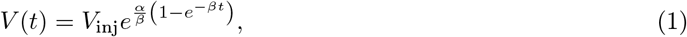

where *V*_inj_ is the initial tumor size at *t*_inj_ = 0 and *α* and *β* are two parameters.

While the etiology of the Gompertz model has been long debated [9], several independent studies have reported a strong and significant correlation between the parameters *α* and *β* in either experimental systems [6, 10, 11], or human data [11, 12, 13]. While some authors suggested this would imply a constant maximal tumor size (given by 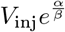 in (1)) across tumor types within a given species [11], others argued that because of the presence of the exponential function, this so called ‘carrying capacity’ could vary over several orders of magnitude [14]. To date, the generalizability, implications and understanding of this observation are still a source of active debate in the oncology modeling community.

Mathematical models for tumor growth have been previously studied and compared at the level of individual kinetics and for prediction of future tumor growth [15, 16]. However, to our knowledge, a detailed study of statistical properties of classical growth models at the level of the population (i.e. integrating structural dynamics with inter-animal variability) yet remains to be reported. Longitudinal data analysis with nonlinear mixed-effects is an ideal tool to perform such a task [17, 18]. In addition, the reduced number of parameters (from *p* × *N* to 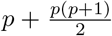 where *N* is the number of animals and *p* the number of parameters of the model) ensures higher robustness (smaller standard errors) of the estimates. This framework is particularly adapted to study the above-mentioned correlation of the Gompertz parameter estimates.

Moreover, using population distributions as priors allows to make predictions on new subjects by means of Bayesian algorithms [19, 20, 21]. The added value of the latter methods is that only few measurements per individual are necessary to obtain reliable predictions. In contrast with previous work focusing on the *forward* prediction of the size of a tumor [15], the present study focuses on the *backward* problem, i.e. the estimation of the age of a tumor [22]. This question is of fundamental importance in the clinic since the age of a tumor can be used as a proxy for determination of the invisible metastatic burden at diagnosis [23]. In turn, this estimation has critical implications for decision of the extent of adjuvant therapy [24]. Since predictions of the initiation time of clinical tumors are hardly possible to verify for clinical cases, we developed and validated our method using experimental data from multiple data sets in several animal models. This setting allowed to have enough measurements, on a large enough time frame in order to assess the predictive power of the methods.

## 2 Material and methods

### 2.1 Mice experiments

The experimental data comprised three data sets. Animal tumor model studies were performed in strict accordance with guidelines for animal welfare in experimental oncology and were approved by local ethics committees. Precise description of experimental protocols was reported elsewhere (see [15] for the volume measurements and [25] for the fluorescence measurements).

#### Breast data measured by volume (N = 66)

This data consisted of human LM2-4LUC+ triple negative breast carcinoma cells originally derived from MDA-MB-231 cells. Animal studies were performed as described previously under Roswell Park Comprehensive Cancer Center (RPCCC) Institutional Animal Care and Use Committee (IACUC) protocol number 1227M [PMID: 25167199 and 26511632]. Briefly, animals were orthotopically implanted with LM2-4LUC+ cells (106 cells at injection) into the right inguinal mammary fat pads of 6- to 8-week-old female severe combined immunodeficient (SCID) mice. Tumor size was measured regularly with calipers to a maximum volume of 2 cm^3^, calculated by the formula *V* = *π*/6*w*2*L* (ellipsoid) where *L* is the largest and *w* is the smallest tumor diameter. The data was pooled from eight experiments conducted with a total of 581 observations. All LM2-4LUC+ implanted animals used in this study are vehicle-treated animals from published studies [PMID: 25167199 and 26511632]. Vehicle formulation was carboxymethylcellulose sodium (USP, 0.5% w/v), NaCl (USP, 1.8% w/v), Tween-80 (NF, 0.4% w/v), benzyl alcohol (NF, 0.9% w/v), and reverse osmosis deionized water (added to final volume) and adjusted to pH 6 (see [PMID: 18199548]) and was given at 10ml/kg/day for 7-14 days prior tumor resection.

#### Breast data measured by fluorescence (N = 8)

This data consisted of human MDA-MB-231 cells stably transfected with dTomato lentivirus. Animals were orthotopically implanted (80,000 cells at injection) into the mammary fat pads of 6-week-old female nude mice. Tumor size was monitored regularly with fluorescence imaging. The data comprised a total of 64 observations. To recover the fluorescence value corresponding to the injected cells, we computed the ratio between the fluorescence signal and the volume measured in mm^3^. We used linear regression considering the volume data of a different data set with same experimental setup (mice, tumor type and number of injected cells). The estimated ratio was 1.52· 10^9^ photons/(s·mm^3^) with relative standard error of 11.3%, therefore the initial fluorescence signal was 1.22· 10^7^ photons/s.

#### Lung data measured by volume (N = 20)

This data consisted of murine Lewis lung carcinoma cells originally derived from a spontaneous tumor in a C57BL/6 mouse [26]. Animals were implanted subcutaneously (10^6^ cells at injection) on the caudal half of the back in anesthetized 6- to 8-week-old C57BL/6 mice. Tumor size was measured as described for the breast data to a maximum volume of 1.5 cm^3^. The data was pooled from two experiments with a total of 188 observations.

### 2.2 Tumor growth models

We denoted by *t_I_* and *V_I_* the initial conditions of the equation. At time of injection (*t*_inj_ = 0), we assumed that all tumor volumes within a group had the same volume *V*_inj_ (taken to be equal to the number of injected converted in the appropriate unit) and denoted by *α* the specific growth rate (i.e. 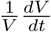) at this time and volume.

We considered the Exponential, Logistic and Gompertz models [15]. The first two are respectively defined by the following equations

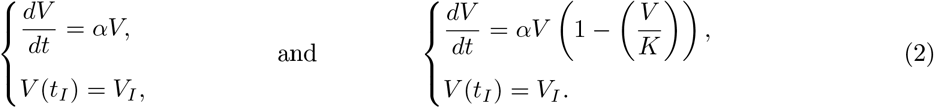

In the Logistic equation, *K* is a carrying capacity parameter. It expresses a maximal reachable size due to competition between the cells (e.g. for space or nutrients).

The Gompertz model is characterized by an exponential decrease of the specific growth rate with rate denoted here by *β*. Although multiple expressions and parameterizations coexist in the litterature, the definition we adopted here reads as follows:

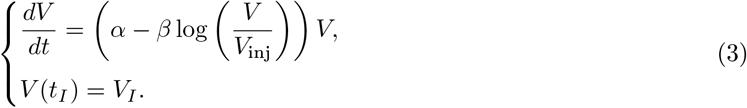

Note that the injected volume *V*_inj_ appears in the differential equation defining *V*. This is a natural consequence of our assumption of *α* as being the specific growth rate at *V* = *V*_inj_. This model exhibits sigmoidal growth up to a saturating value given by 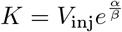.

### 2.3 Population approach

Let *N* be the number of subjects within a population (group) and 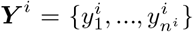 the vector of longitudinal measurements in animal *i*, where 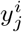 is the observation of subject *i* at time 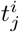 for *i* = 1, …, *N* and *j* = 1, …, *n^i^* (*n^i^* is the number of measurements of individual *i*). We assumed the following observation model

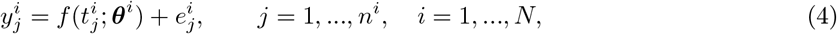

where 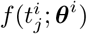 is the evaluation of the tumor growth model at time 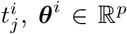 is the vector of the parameters relative to the individual *i* and 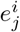 the residual error model, to be defined later. An individual parameter vector ***θ***^*i*^ depends on fixed effects ***μ***, identical within the population, and on a random effect ***η***^*i*^, specific to each animal. Random effects follow a normal distribution with mean zero and variance matrix ***ω***. Specifically:

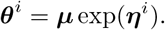

We considered a combined residual error model 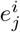, defined as

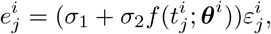

where 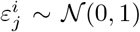 are the residual errors and ***σ*** = [*σ*_1_, *σ*_2_] is the vector of the residual error model parameters.

In order to compute the population parameters, we maximized the population likelihood, obtained by pooling all the data together. Usually, this likelihood cannot be computed explicitely for nonlinear mixed-effect models. We used the stochastic approximation expectation minimization algorithm (SAEM) [17], implemented in the `Monolix 2018 R2` software [27].

In the remainder of the manuscript we will denote by *ϕ* = {***μ***, ***ω***, ***σ***} the set of the population parameters containing the fixed effects ***μ***, the covariance of the random effects ***ω*** and the error model parameters ***σ***.

### 2.4 Individual predictions

For a given animal *i*, the backward prediction problem we considered was to predict the age of the tumor based on the three last measurements 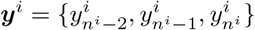. Since we were in an experimental setting, we considered the injection time as the initiation time and thus the age was given by 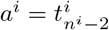. To avoid using knowledge from the past, we first shifted the times of measurement by 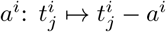. Then, we considered as model *f*(*t*; *θ^i^*) the solution of the Cauchy problem (3) endowed with initial conditions 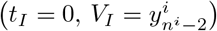. For estimation of the parameters (estimate 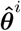), we applied two different methods: likelihood maximization alone (no use of prior population information) and Bayesian inference (use of prior). The predicted age 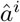 was then defined by

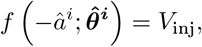

that is:

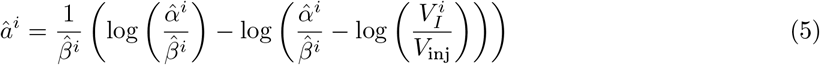

in case of the Gompertz model.

#### 2.4.1 Likelihood maximization

For individual predictions with likelihood maximization, no prior information on the distribution of the parameters was used. Parameters of the error model were not re-estimated: values from the population analysis were used. The constant part *σ*_1_ was found negligible compared to the large volumes at late times, thus only the proportional term of the error model was used (*σ* = *σ*_2_). The log-likelihood can be derived from (4):

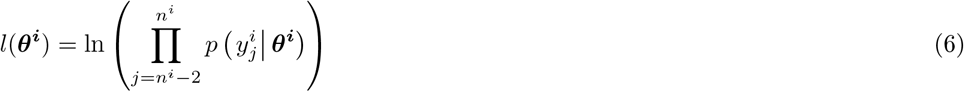

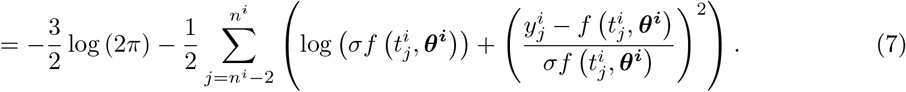

In order to guarantee the positivity of the parameters, we introduced the relation ***θ***^*i*^ = *g*(***γ***^*i*^) = *e*^*γ*^*i*^^ and substituted this in equation(7). The negative of equation (7) was minimized with respect to ***γ^i^*** (yielding the maximum likelihood estimate 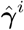) with the function `minimize` of the python module `scipy.optimize`, for which the Nelder-Mead algorithm was applied. Thanks to the invariance property, the maximum likelihood estimator of ***θ***^*i*^ was determined as 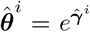. Individual prediction intervals were computed by sampling the parameters ***θ***^*i*^ from a gaussian distribution with variance-covariance matrix of the estimate defined as 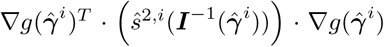 where 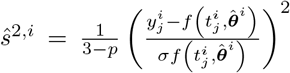, with *p* the number of parameters and 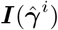 and 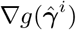 the Fisher information matrix and the gradient of the function *g*(***γ***) evaluated, respectively, in the estimate 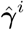. Denoting by 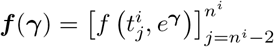 and by 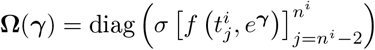, the Fisher information matrix was defined by [28]:

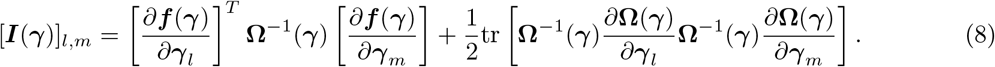

#### 2.4.2 Bayesian inference

When applying the Bayesian method, we considered *training sets* to learn the distribution of the parameters *ϕ* and *test sets* to derive individual predictions. For a given animal *i* of a *test set*, we predicted the age of the tumor based on the combination of: 1) population parameters *ϕ* identified on the *training set* using the population approach and 2) the three last measurements of animal *i*. We set as initial conditions *t_I_* = 0 and 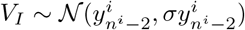. This last assumption was made to account for measurement uncertainty on 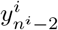. We then estimated the posterior distribution *p*(***θ***^*i*^|***y***^*i*^) of the parameters ***θ***^*i*^ using a Bayesian approach [20]:

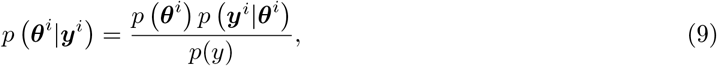

where *p*(***θ***^*i*^) is the prior distribution of the parameters estimated through nonlinear mixed-effects modeling and *p*(***y***^*i*^|***θ***^*i*^) is the likelihood, defined from equation (4). The predicted distributions of extrapolated growth curves and subsequent 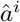 were computed by sampling ***θ***^*i*^ from its posterior distribution (9) using `Pystan`, a Python interface to the software `Stan` [21] for Bayesian inference based on the No-U-Turn sampler, a variant of Hamiltonian Monte Carlo [19]. Predictions of 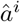 were then obtained from (5), considering the median value of the distribution.

Different data sets were used for learning the priors (*training sets*) and prediction (*test sets*) by means of *k*-fold cross validation, with *k* equal to the total number of animals of the dataset (*k* = *N*, i.e. leave-one-out strategy). At each iteration we computed the parameters distribution of the population composed by *N* – 1 individuals and used this as prior to predict the initiation time of the excluded subject *i*. The `Stan` software was used to draw 2000 realizations from the posterior distribution of the parameters of the individual *i*.

## 3 Results

The results reported below were similar for the three data sets presented in the materials and methods. For conciseness, the results presented herein are related to the large dataset (breast cancer data measured by volume). Results relative to the other (smaller) datasets are reported in the Supplemental.

### 3.1 Population analysis of tumor growth curves

The population approach was applied to test the descriptive power of the Exponential, Logistic and Gompertz models for tumor growth kinetics. The number of injected cells at time *t*_inj_ = 0 was 10^6^, therefore we fixed the initial volume *V*_inj_ = 1 mm^3^ in the whole dataset [15]. We set (*t_I_*, *V_I_*) = (*t*_inj_, *V*_inj_) as initial condition of the equations.

We ran the SAEM algorithm with the `Monolix` software to estimate the fixed and random effects [27]. Moreover, we evaluated different statistical indices in order to compare the different tumor growth models. This also allowed learning of the parameter population distributions that were used later as priors for individual predictions. Results are reported in Table 1, where the models are ranked according to their AIC (Akaike Information Criterion), a metrics combining parsimony and goodness-of-fit. The Gompertz model was the one with the lowest values, indicating superior goodness-of-fit. This was confirmed by diagnostic plots (Figure 1). The visual predictive checks (VPCs) in Figure 1A compare the empirical percentiles with the theoretical percentiles, i.e. those obtained from simulations of the calibrated models. The VPC of the Exponential and Logistic models showed clear model misspecification. On the other hand, the VPC of the Gompertz model was excellent, with observed percentiles close to the predicted ones and small prediction intervals (indicative of correct identifiability of the parameters). Figure 1B shows the prediction distribution of the three models. This allowed to compare the observations with the theoretical distribution of the predictions. Only the prediction distribution of the Gompertz model covered the entire dataset. The Logistic model exhibited a saturation of tumor dynamics at lower values than compatible with the data.

**Table 1.**
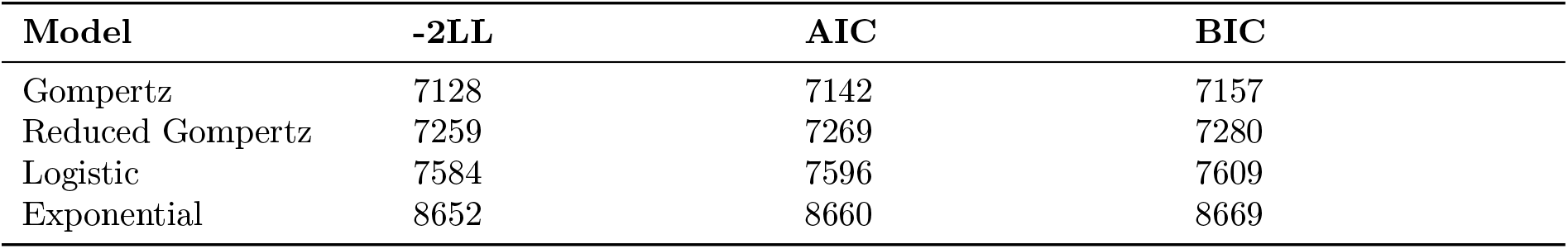
Models ranked in ascending order of AIC (Akaike information criterion). Other statistical indices are the log-likelihood estimate (−2LL) and the Bayesian information criterion (BIC).

| Model | -2LL | AIC | BIC |
| --- | --- | --- | --- |
| Gompertz | 7128 | 7142 | 7157 |
| Reduced Gompertz | 7259 | 7269 | 7280 |
| Logistic | 7584 | 7596 | 7609 |
| Exponential | 8652 | 8660 | 8669 |

**Figure 1.**
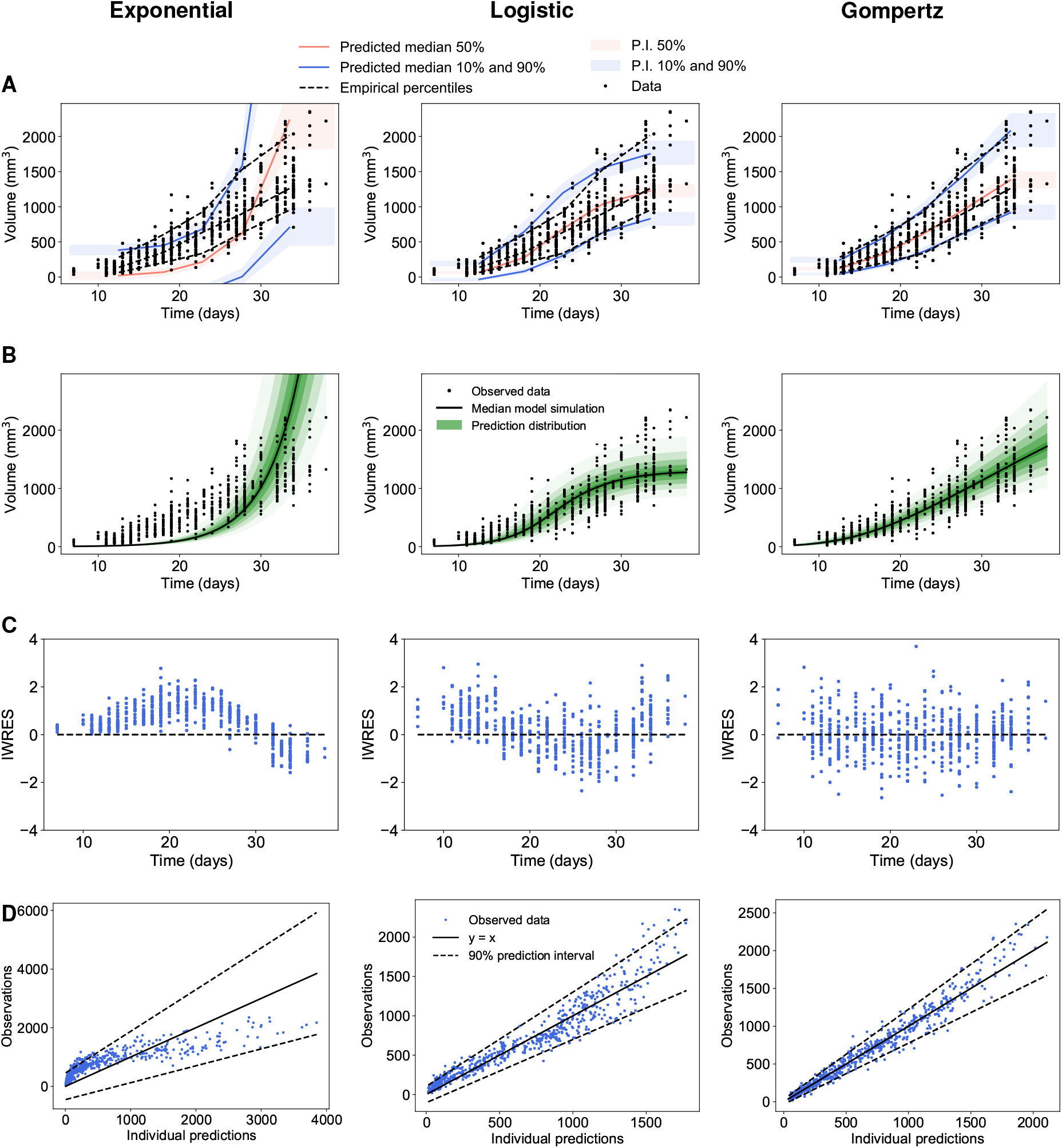
Population analysis of experimental tumor growth kinetics. A) Visual predictive checks assess goodness-of-fit for both structural dynamics and inter-animal variability by reporting model-predicted percentiles (together with confidence prediction intervals (P.I) in comparison to empirical ones. B) Prediction distributions. C) Individual weighted residuals (IWRES) with respect to time. D) Observations vs predictions Left: Exponential, Center: Logistic, Right: Gompertz models.

Moreover, the distribution of the residuals was symmetrical around a mean value of zero with the Gompertz model (Figure 1C), strengthening its good descriptive power, while the Exponential and Logistic models exhibited clear skewed distributions. The observations vs individual predictions in Figure 1D further confirmed these findings.

These observations at the population level were confirmed by individual fits, computed from the mode of the posterior conditional parameter distribution for each individual (Figure 2). Confirming previous results [15], the optimal fits of the Exponential and Logistic models were unable to give appropriate description of the data, suggesting that these models should not be used to describe tumor growth, at least in similar settings to ours. Fitting of late timepoints data forced the proliferation parameter of the Exponential model to converge towards a rather low estimate, preventing reliable description of the early datapoints. The converse occurred for the Logistic. Constrained by the early data points imposing to the model the pace of the growth deceleration, the resulting estimation of the carrying capacity *K* was biologically irrelevant (much too small, typical value 1332 mm^3^, see Table 2), preventing the model to give a good description of the late growth.

**Figure 2.**
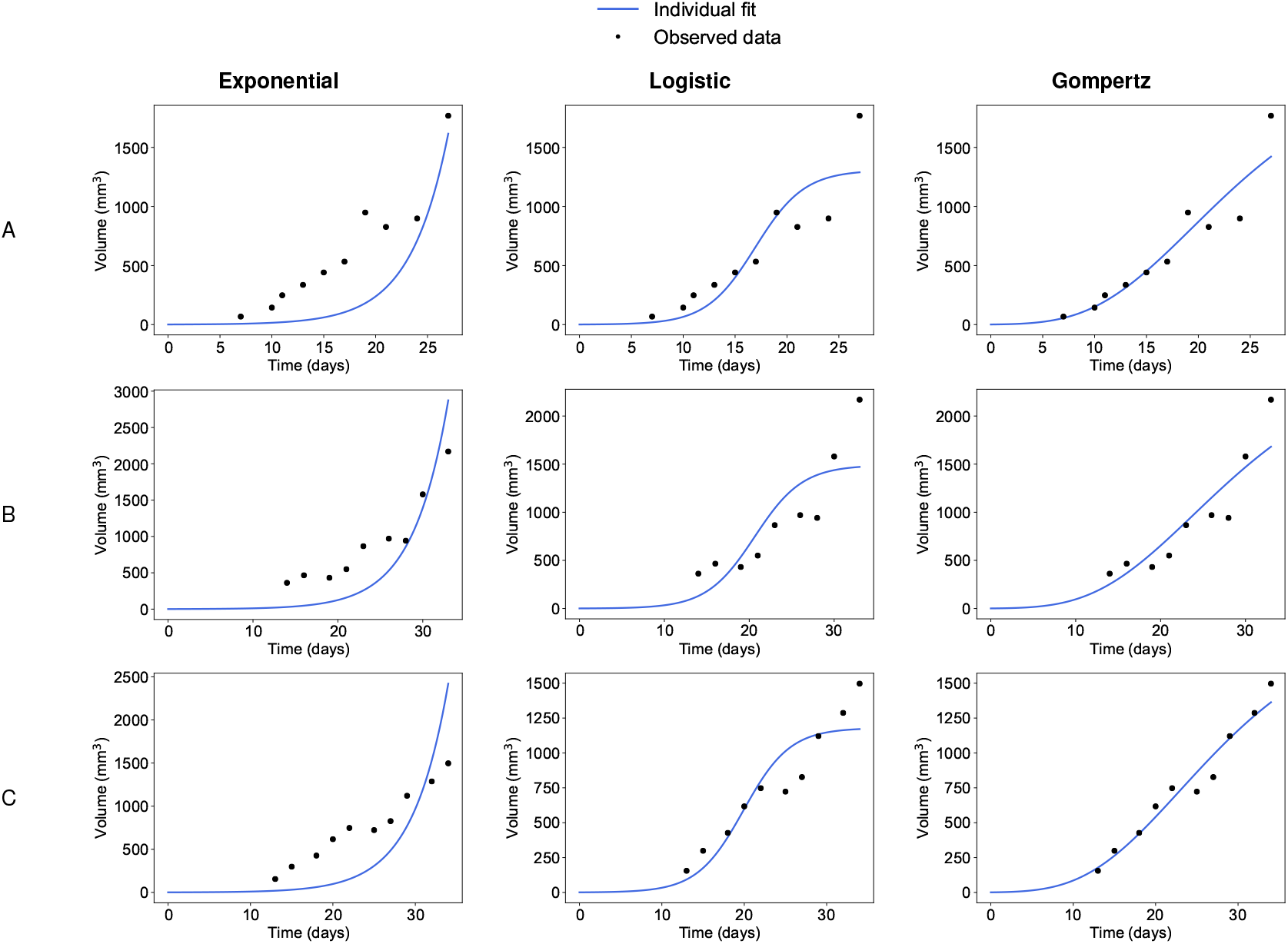
Individual fits from population analysis. Three representative examples of individual fits computed with the population approach relative to the Exponential (left), the Logistic (center) and the Gompertz (right) models.

**Table 2.** Fixed effects (typical values) of the parameters of the different models. CV = Coefficient of Variation, expressed in percentage and estimated as the standard deviation of the parameter divided by the fixed effect and multiplied by 100. ***σ*** is vector of the residual error model parameters. Last column shows the relative standard errors (R.S.E.) of the estimates.

| Model | Parameter | Unit | Fixed effects | CV (%) | R.S.E. (%) |
| --- | --- | --- | --- | --- | --- |
| Gompertz | $\alpha$ | day<sup>-1</sup> | 0.573 | 34.73 | 2.56 |
| | $\beta$ | day<sup>-1</sup> | 0.0705 | 391.49 | 3.61 |
| | $\sigma$ | - | [19.1, 0.12] | | [18.3, 7.36] |
| Reduced Gompertz | $\beta$ | day<sup>-1</sup> | 0.0725 | 180.69 | 1.91 |
| | $k$ | - | 7.98 | 0 | 0.363 |
| | $\sigma$ | - | [13.9, 0.183] | | [22.3, 5.17] |
| Logistic | $\alpha$ | day<sup>-1</sup> | 0.324 | 42.90 | 1.88 |
| | $K$ | mm<sup>3</sup> | 1332 | 0.02 | 4.39 |
| | $\sigma$ | - | [57.2, 0.136] | | [9.8, 8.74] |
| Exponential | $\alpha$ | day<sup>-1</sup> | 0.229 | 34.98 | 1.35 |
| | $\sigma$ | - | [283, 0.254] | | [6.06, 14.3] |

Table 2 provides the values of the population parameters. The relative standard error estimates associated to population parameters were all rather low (<4.39%), indicating good practical identifiability of the model parameters. Standard error estimates of the constant error model parameters were found to be slightly larger (<22.3%), suggesting that for some models a proportional error model might have been more appropriate - but not in case of the Exponential model. Since our aim was to compare different tumor growth equations, we established a common error model parameter, i.e. a combined error model. Relative standard errors of the standard deviations of the random effects ***ω*** were all smaller than 9.6% (not shown).

These model findings in the breast cancer cell line were further validated with the other cell lines. For both the lung cancer and the fluorescence-breast cancer cell lines, the Gompertz model outperformed the other competing models (see Supplementary Tables S1 and Tables S2 for the two data sets), as also shown by the diagnostic plots (Figures S1,S2). For the fluorescence-breast cancer cell line we used a proportional error model (i.e., we fixed *σ*_1_ = 0). In this case the inter-individual variability was found to be modest. This was due to the small number of animals in the data set and to a considerable intra-individual variability (Supplementary Figure S4) associated to large measurement error (see Table S4).

Together, these results confirmed that the Exponential and Logistic models are not appropriate models of tumor growth while the Gompertz model has excellent descriptive properties, for both goodness-of-fit and parameter identifiability purposes.

### 3.2 The reduced Gompertz model

#### 3.2.1 Correlation between the Gompertz parameters

During the estimation process of the Gompertz parameters, we found a high correlation between *α* and *β* within the population. At the population level, the SAEM algorithm estimated a correlation of the random effects equal to 0.981. At the individual level, *α^i^* and *β^i^* were also highly linearly correlated (Figure 3A, *R*^2^ = 0.968). This motivated the reformulation of the alpha parameter as follows:

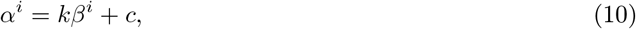

where *k* and *c* are representing the slope and the intercept of the regression line, respectively. In our analysis we found *c* to be small (*c* = 0.14), thus we further assumed this term to be negligible and fixed it to 0. This suggests *k* as a characteristic constant of tumor growth within a given animal model [11, 29].

**Figure 3.**
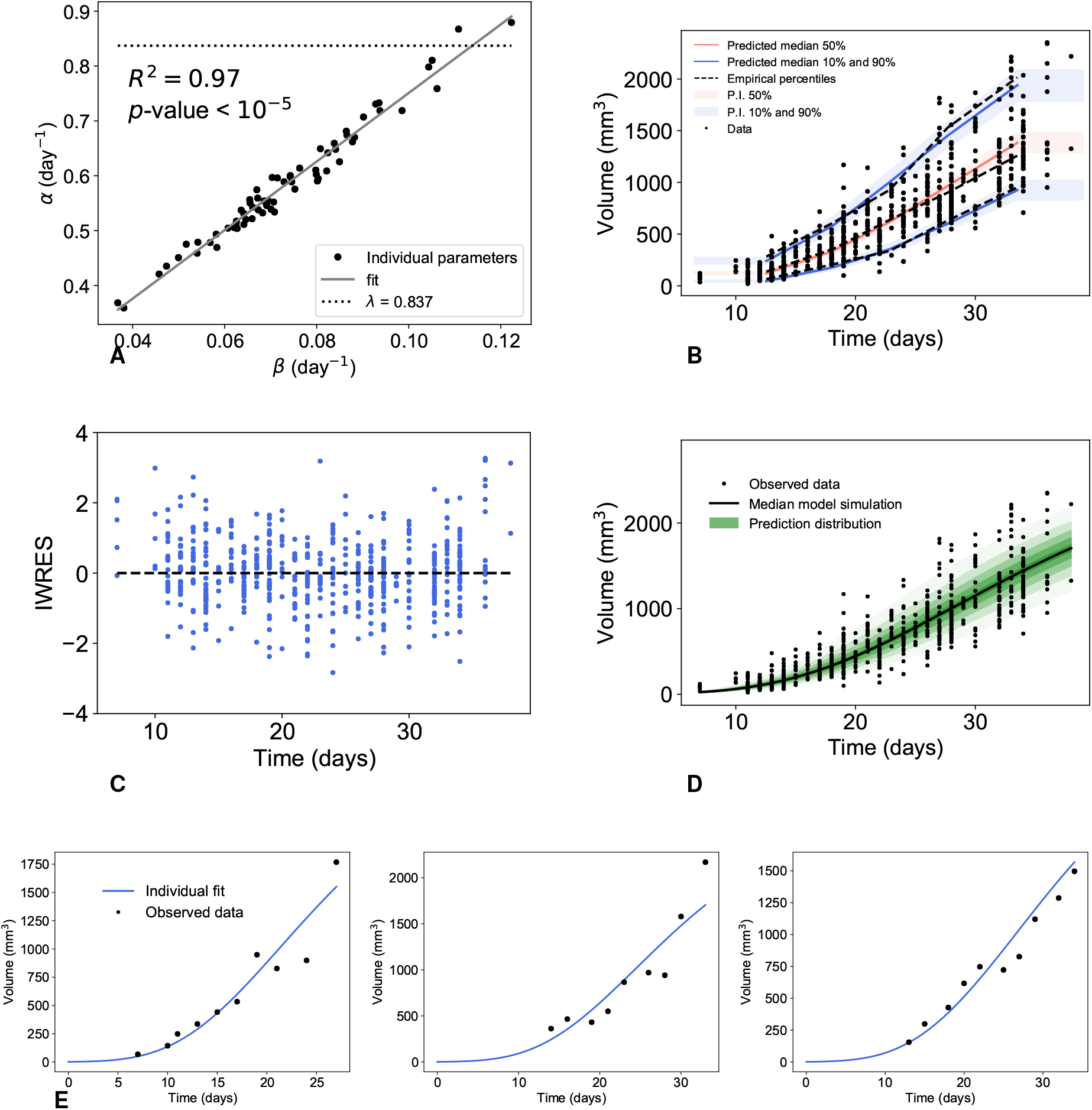
Correlation of the Gompertz parameters and diagnostic plots of the reduced Gompertz model from population analysis. Correlation between the individual parameters of the Gompertz model (A) and results of the population analysis of the reduced Gompertz model: visual predictive check (B), examples of individual fits (C) and scatter plots of the residuals (D).

In turn, this implies an approximately constant limiting size

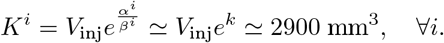

The other data sets gave analogous results. The estimated correlations of the random effects were 0.967 and 0.998 for the lung cancer and for the fluorescence-breast cancer, respectively. The correlation between the parameters was also confirmed at the individual level (see Supplementary Figures S5A and Supplementary Figures S6A, R^2^ was 0.961 and 0.99 for the two data sets, respectively).

#### 3.2.2 Biological interpretation in terms of the proliferation rate

By definition, the parameter α is the specific growth rate at the time of injection. Assuming that the cells don’t change their proliferation kinetics when implanted, this value should thus be equal to the *in vitro* proliferation rate (supposed to be the same for all the cells of the same cell line), denoted here by λ. The value of this biological parameter was assessed *in vitro* and estimated at 0.837 [23]. Confirming our theory, we indeed found estimated values of α close to λ (fixed effects of 0.585), although strictly smaller estimates were reported in the majority of cases (Figure 3A). We postulated that this difference could be explained by the fact that not all the cells will be successfully grafted when injected in an animal. Denoting by 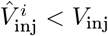 the volume of these cells, our mathematical expression of λ would now read as:

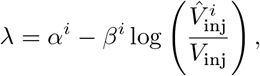

which is consistent with our findings since this leads to values of λ > *α^i^*. In turn, this gives estimates of the percentage of successful egraftment at 18% ± 5.9%.

#### 3.2.3 Population analysis of the reduced Gompertz model

The high correlation among the Gompertz parameters, combined to the biological rationale explained above, suggested that a reduction of the degrees of freedom (number of parameters) in the Gompertz model could improve identifiability and yield a more parsimonious model. We considered the expression (10), assuming c negligible. We therefore propose the following reduced Gompertz model:

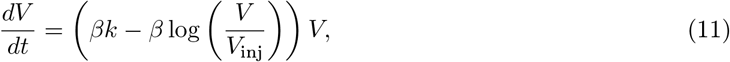

where *β* has mixed effects, while *k* has only fixed effects, i.e., is constant within the population.

Figure 3 shows the results relative to the population analysis performed. Results of the diagnostic plots indicated no deterioration of the goodness-of-fit as compared with the Gompertz model (Figure 3B-D). Only on the last timepoint was the model slightly underestimating the data (Figure 3D), which might explain why the model performs slightly worse than the two-parameters Gompertz model in terms of strictly quantitative statistical indices (but still better than the Logistic or Exponential models, Table 1). Individual dynamics were also accurately described (Figure 3E). Parameter identifiability was also excellent (Table 2).

The other two data sets gave similar results (see Supplementary Figures S5 and S6).

Together, these results demonstrated the accuracy of the reduced Gompertz model, with improved robustness as compared to previous models.

### 3.3 Prediction of the age of a tumor

Considering the increased robustness of the reduced Gompertz model (one individual parameter less than the Gompertz model), we further investigated its potential for improvement of predictive power. We considered the problem of estimating the age of a tumor, that is, the time elapsed between initiation (here the time of injection) and detection occurring at larger tumor size (Figure 4). For a given animal *i*, we considered as first observation 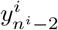 and aimed to predict its age 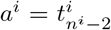 (see Methods). We compared the results given by the Bayesian inference with the ones computed with standard likelihood maximization method (see Methods). To that end, we did not consider any information on the distribution of the parameters. For the reduced Gompertz model however (likelihood maximization case), we used the value of *k* calculated in the previous section (Table 2), thus using information on the entire population. Importantly, for both prediction approaches, our methods allowed not only to generate a prediction of *a^i^* for estimation of the model accuracy (i.e. absolute relative error of prediction), but also to estimate the uncertainty of the predictions (i.e. precision, measured by the width of the 90% prediction interval (PI)).

**Figure 4.**
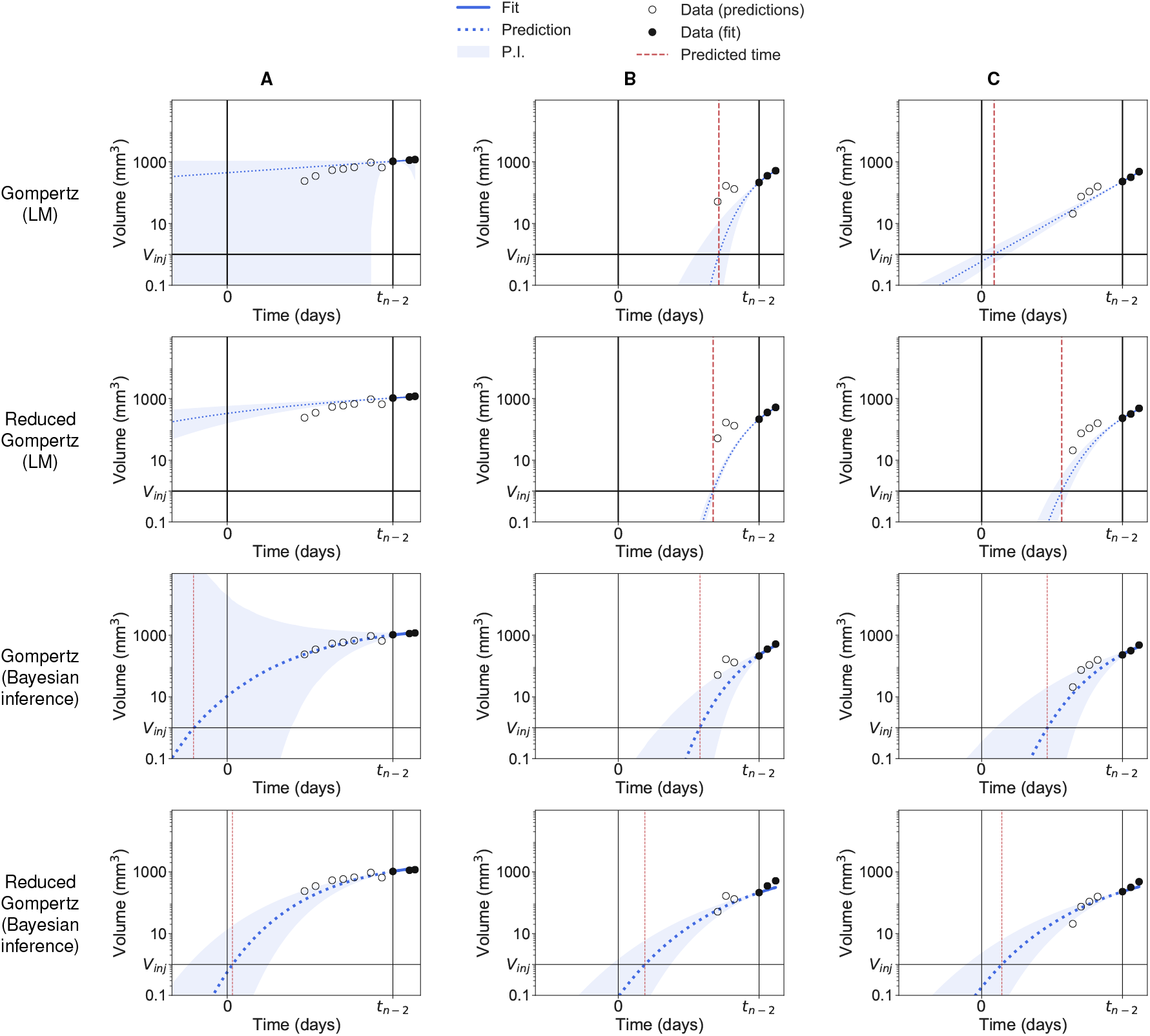
Backward predictions computed with likelihood maximization and with Bayesian inference. Three examples of backward predictions of individuals A, B and C computed with likelihood maximization (LM) and Bayesian inference: Gompertz model with likelihood maximization (first row); reduced Gompertz with likelihood maximization (second row); Gompertz with Bayesian inference (third row) and reduced Gompertz with Bayesian inference (fourth row). Only the last three points are considered to estimate the parameters. The grey area is the 90% prediction interval (P.I) and the dotted blue line is the median of the posterior predictive distribution. The red line is the predicted initiation time and the black vertical line the actual initiation time.

Figure 4 presents a few examples of prediction of three individuals without (LM) and with (Bayesian inference) priors relative to the breast cancer measured by volume. For the other two cell lines, see the Supplementary Figures S7 and S9. The reduced Gompertz model combined to Bayesian inference (bottom row) was found to have the best accuracy in predicting the initiation time (mean error = 12.1%, 9.4% and 12.3% for the volume-breast cancer, lung cancer and fluorescence-breast cancer respectively) and to have the smallest uncertainty (precision = 15.2, 7.34 and 23.6 days for the three data sets, respectively). Table 3 gathers results of accuracy and precision for the Gompertz and reduced Gompertz models under LM and Bayesian inference relative to the three data sets. With only local information of the three last data points, the Gompertz model predictions were very inaccurate (mean error = 205%, 175% and 236%) and the Fisher information matrix was often singular, preventing standard errors to be adequately computed. With one degree of freedom less, the reduced Gompertz model had better performances with LM estimation but still large uncertainty (mean precision under LM = 186, 81.6 and 368 days) and poor accuracy using LM (mean error = 74.1%, 66.1% and 91.7%). Examples shown in Figure 4 were representative of the entire population relative to the breast cancer measured by volume. Eventually, for 97%, 95% and 87.5% of the individuals of the three data sets the actual value of the age fell in the respective prediction interval when Bayesian inference was applied in combination with the reduced Gompertz models. This means a good coverage of the prediction interval and indicates that our precision estimates were correct. On the other hand, this observation was not valid in case of likelihood maximization, where the actual value fell in the respective prediction interval for only 40.9%, 30% and 87.5% of the animals when the reduced Gompertz model was used.

**Table 3.**
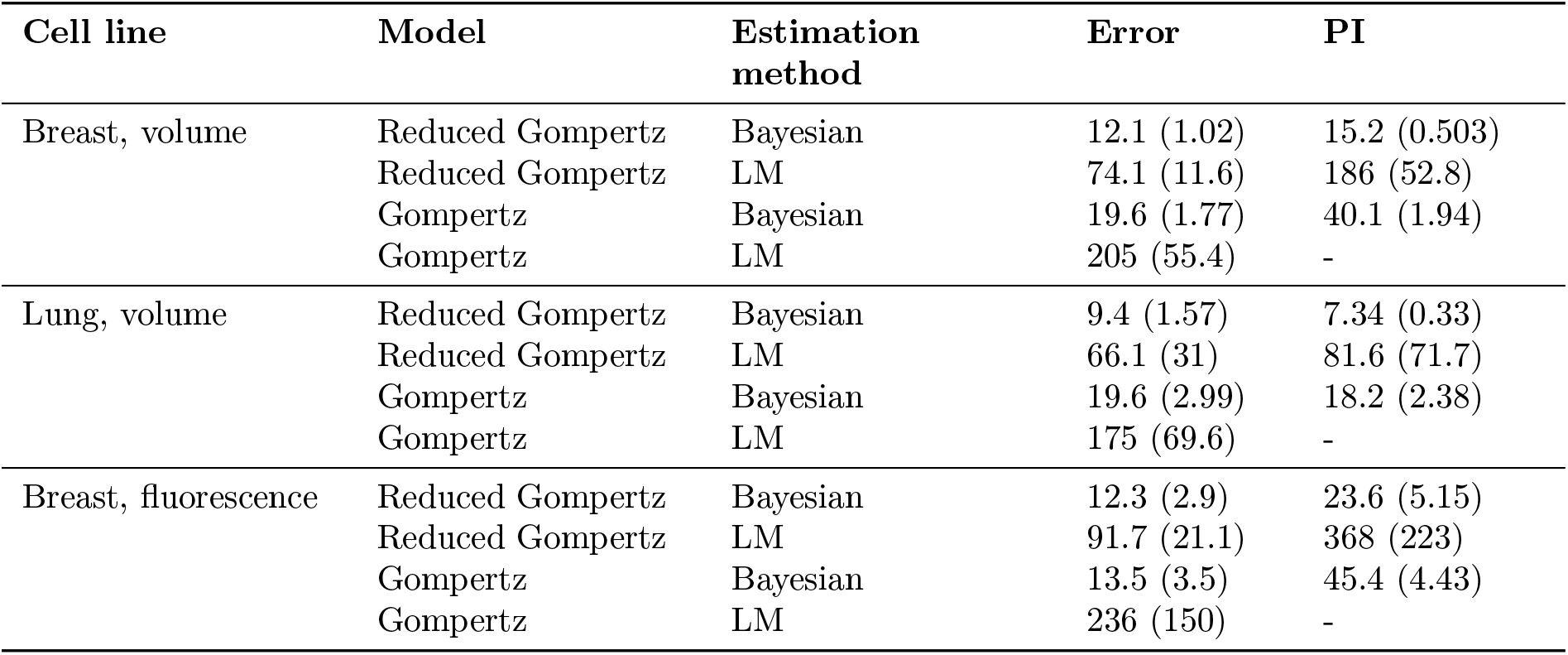
Accuracy and precision of methods for prediction of the age of experimental tumors of the three cell lines. Accuracy was defined as the absolute value of the relative error (in percent). Precision was defined as the width of the 95% prediction interval (in days). Reported are the means and standard errors (in parenthesis). LM = likelihood maximization

| Cell line | Model | Estimation method | Error | PI |
| --- | --- | --- | --- | --- |
| Breast, volume | Reduced Gompertz | Bayesian | 12.1 (1.02) | 15.2 (0.503) |
|  | Reduced Gompertz | LM | 74.1 (11.6) | 186 (52.8) |
|  | Gompertz | Bayesian | 19.6 (1.77) | 40.1 (1.94) |
|  | Gompertz | LM | 205 (55.4) | - |
| Lung, volume | Reduced Gompertz | Bayesian | 9.4 (1.57) | 7.34 (0.33) |
|  | Reduced Gompertz | LM | 66.1 (31) | 81.6 (71.7) |
|  | Gompertz | Bayesian | 19.6 (2.99) | 18.2 (2.38) |
|  | Gompertz | LM | 175 (69.6) | - |
| Breast, fluorescence | Reduced Gompertz | Bayesian | 12.3 (2.9) | 23.6 (5.15) |
|  | Reduced Gompertz | LM | 91.7 (21.1) | 368 (223) |
|  | Gompertz | Bayesian | 13.5 (3.5) | 45.4 (4.43) |
|  | Gompertz | LM | 236 (150) | - |

As a general result, addition of *a priori* population information by means of Bayesian estimation resulted in drastic improvement of the prediction performances (Figure 5). Results relative to the other data sets are shown in Supplementary Figures S7, S8 for the lung cell line and S9, S10 for the breast cell line measured by fluorescence. For the fluorescence-breast cancer cell line we could not report a significant difference in terms of accuracy between the Gompertz and the reduced Gompertz when applying Bayesian inference. This can be explained by the low number of individuals included in the data set.

**Figure 5.**
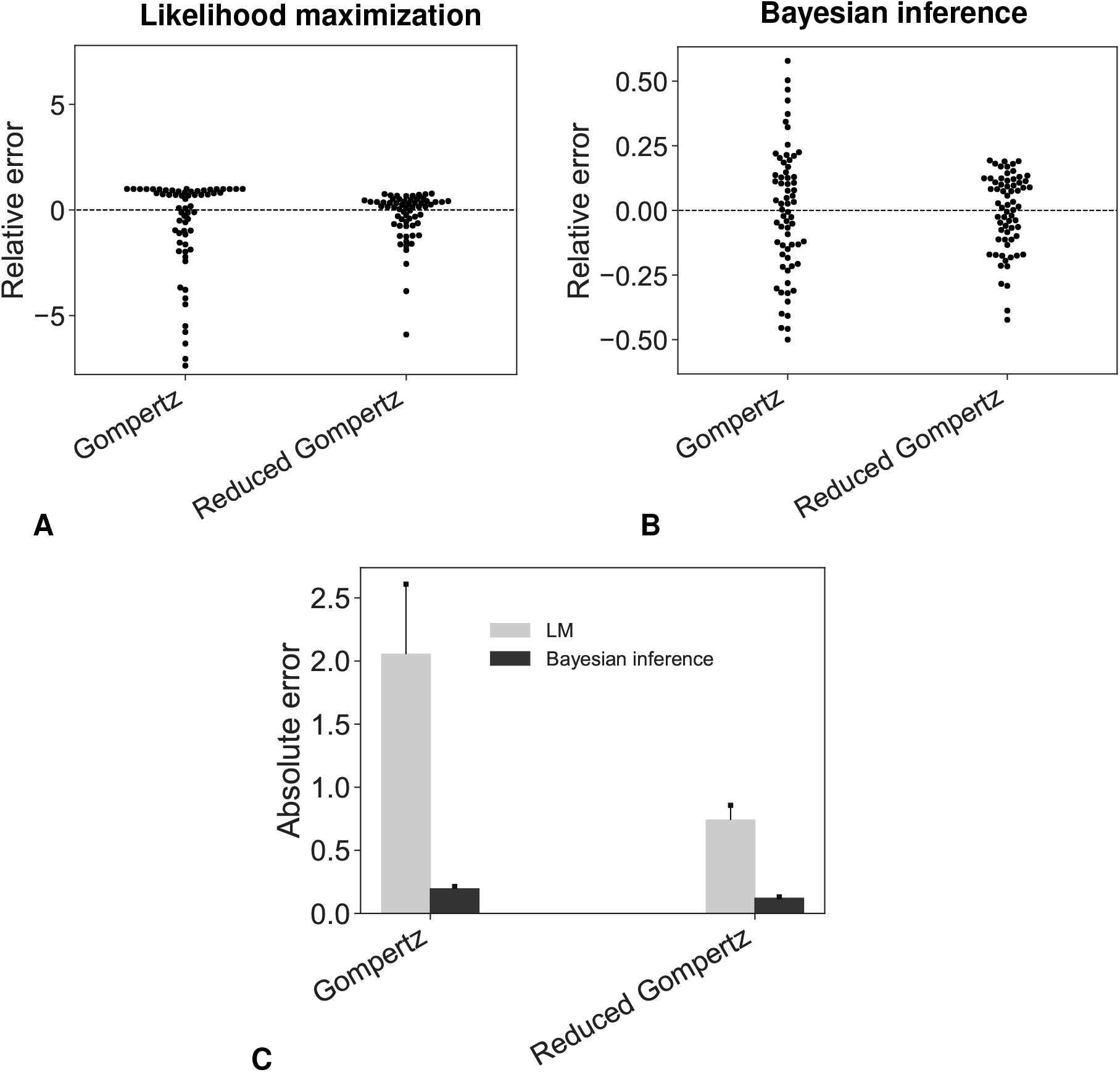
Accuracy of the prediction models. Swarmplots of relative errors obtained under likelihood maximization (A) or Bayesian inference (B). (C) Absolute errors. In (A) four extreme outliers were omitted (values of the relative error were greater than 20) for both the Gompertz and the reduced Gompertz in order to ensure readability.

Overall, the combination of the reduced Gompertz model with Bayesian inference clearly outperformed the other methods for prediction of the age of experimental tumors.

## 4 Discussion

We have analyzed tumor growth curves from multiple animal models and experimental techniques, using a population framework (nonlinear mixed-effects [17]). This approach is ideally suited for experimental or clinical data of the same tumor types within a given group of subjects. Indeed, it allows for a description of the inter-subject variability that is impossible to obtain when fitting models to averaged data (as often done for tumor growth kinetics [30]), while enabling robust population-level description that does not require individual fits. As expected from the classical observation of decreasing specific growth rates [6, 31, 8, 32, 33], the Exponential model generated very poor fits. More surprisingly given its popularity in the theoretical community (probably due to its ecological ground), the Logistic model was also rejected, due to unrealistically small inferred value of the carrying capacity *K*. This finding confirms at the population level previous results obtained from individual fits [15, 34]. It suggests that the underlying theory (competition between the tumor cells for space or nutrients) is unable – at least when considered alone – to explain the d decrease of the specific growth rate, suggesting that additional mechanisms need to be accounted for. Few studies have previously compared the descriptive performances of growth models on the same data sets [15, 35, 16]. In contrast to our results, Vaidya and Alexandro [16] found admissible description of tumor growth data employing the Logistic model. Beyond the difference of animal model, we believe the major reason explaining this discrepancy is the type of error model that was employed, as also noticed by others [34]. Here we used a combined error model, in accordance to our previous study that had examined repeated measurements of tumor size and concluded to rejection of a constant error model (used in [16]). To avoid overfitting, we also made the assumption to keep the initial value *V_I_* fixed to *V*_inj_. As noted before [15], releasing this constraint leads to acceptable fits by either the Exponential or Logistic models (to the price of deteriorated identifiability). However, the estimated values of *V_I_* are in this case are biologically inconsistent.

On the other hand, the Gompertz model demonstrated excellent goodness-of-fit in all the experimental systems that we investigated. This is in agreement with a large body of previous experimental and clinical research work using the Gompertz model to describe unaltered tumor growth in syngeneic [36, 6, 10, 34] and xenograft [37, 38] preclinical models, as well as human data [32, 13, 12, 8]. The poor performances of the Logistic model compared to the Gompertz model can be related to the structural properties of the models. The two sigmoid functions lie between two asymptotes (*V* = 0 and *V* = *K*) and are characterized by an initial period of fast growth followed by a phase of decreasing growth. These two phases are symmetrical in the Logistic model, that is indeed characterized by a decrease of the specific growth rate 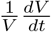 at constant speed. On the other hand, the Gompertz model exhibits a faster decrease of the specific growth rate, at speed 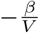, or 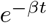 as a function of *t*, and the sigmoidal curve is not symmetric around its inflexion point.

Similarly to previous reports [6, 10, 39, 11, 12, 13], we also found a very strong correlation between the two parameters of the Gompertz model, i.e. *α* the proliferation rate at injection and *β* the rate of decrease of the specific growth rate. Of note, this is not due to a lack of identifiability of the parameters at the individual level, which we investigated and found to be excellent. Such finding motivated our choice to introduce a novel model, the reduced Gompertz model, with only one individual-specific parameter, and one population-specific parameter. We rigorously assessed its descriptive power and found that performances were similar to the two-parameters Gompertz model. Critically, while previous work had demonstrated that two individual parameters were sufficient to describe tumor growth curves [15], these new results now show that this number can be reduced to one. Interestingly, the values of *k* that we inferred were different for the breast and the lung cancer cell lines measured in volume (*k* = 7.98 - 9.51, respectively), in contrast with previous results [11]. This suggests that there might not be a characteristic constant of tumor growth within a species [29] but the correlation could be a typical feature of a tumor type in an animal model. Indeed a small variation of the parameter *k* is associated to a large variation of the carrying capacity *K* = *V*_inj_ exp(*k*). Moreover, we believe that our formulations of the Gompertz (3) and reduced Gompertz (11) give to α a physiological meaning (the *in vitro* proliferation rate) that could be used clinically to predict past or future tumor growth kinetics based on proliferation assays, derived for example from a patient’s tumor sample.

The reduced Gompertz model, combined to Bayesian estimation from the population prior, allowed to reach good levels of accuracy and precision of the time elapsed between the injection of the tumor cells and late measurements, used as an experimental surrogate of the age of a given tumor. Importantly, performances obtained without using a prior were substantially worse. The method proposed herein remains to be extended to clinical data, although it will not be possible to have a firm confirmation since the natural history of neoplasms since their inception cannot be reported in a clinical setting. Nevertheless, the encouraging results obtained here could allow to give approximative estimates. Importantly, the methods we developed also provide a measure of precision, which would give a quantitative assessment of the reliability of the predictions. For clinical translation, Vinj should be replaced by the volume of one cell *V_c_* = 10^−6^ mm^3^. Moreover, because the Gompertz model has a specific growth rate that tends to infinity when V gets arbitrarily small, our results might have to be adapted with the Gomp-Exp model [40, 23].

Personalized estimations of the age of a given patient’s tumor would yield important epidemiological insights and could also be informative in routine clinical practice [22]. By estimating the period at which the cancer initiated, it could give clues on the possible causes (environmental or behavioral) of neoplastic formation. Moreover, reconstruction of the natural history of the pre-diagnosis tumor growth might inform on the presence and extent of invisible metastasis at diagnosis. Indeed, an older tumor has a greater probability of having already spread than a younger one. Altogether, the present findings contribute to the development of personalized computational models of metastasis [23, 41].

## Supplementary Material

### Population analysis

**Table S1.** (lung). Statistical indices of the tumor growth models. **Lung, volume.** Models ranked in ascending order of AIC (Akaike information criterion). Other statistical indices are the log-likelihood estimate (−2LL) and the Bayesian information criterion (BIC).

| Model | -2LL | AIC | BIC |
| --- | --- | --- | --- |
| Gompertz | 2232 | 2246 | 2253 |
| Reduced Gompertz | 2256 | 2266 | 2271 |
| Logistic | 2315 | 2327 | 2333 |
| Exponential | 2644 | 2652 | 2656 |

**Table S2.** (breast-fluorescence). Statistical indices of the tumor growth models. **Breast, fluorescence.** Models ranked in ascending order of AIC (Akaike information criterion). Other statistical indices are the log-likelihood estimate (−2LL) and the Bayesian information criterion (BIC).

| Model | -2LL | AIC | BIC |
| --- | --- | --- | --- |
| Reduced Gompertz | 2953 | 2961 | 2962 |
| Gompertz | 2953 | 2965 | 2966 |
| Logistic | 3009 | 3019 | 3020 |
| Exponential | 3097 | 3103 | 3104 |

**Table S3.**
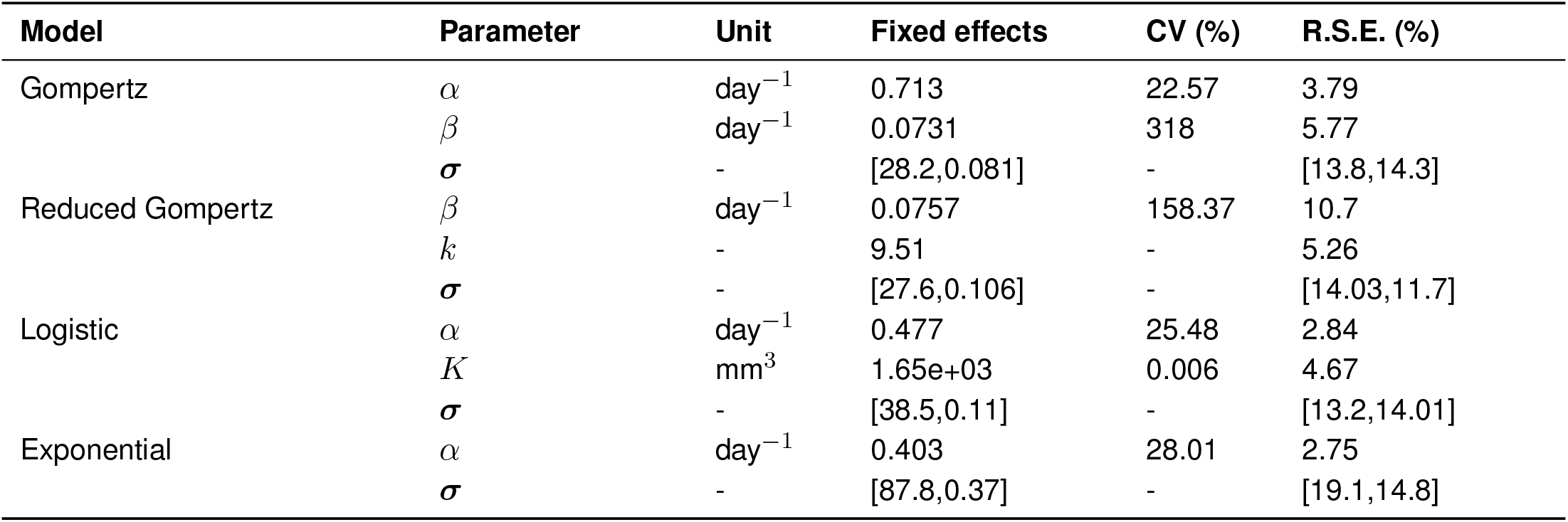
(lung). Parameter values estimated with the SAEM algorithm. **Lung, volume.** Fixed effects (typical values) of the parameters of the different models. CV = Coefficient of Variation, expressed in percentage and estimated as the standard deviation of the parameter divided by the fixed effect and multiplied by 100. ***σ*** is vector of the residual error model parameters. Last column shows the relative standard errors (R.S.E.) of the estimates.

**Table S4.** (breast-fluorescence). Parameter values estimated with the SAEM algorithm. **Breast, fluorescence.** Fixed effects (typical values) of the parameters of the different models. CV = Coefficient of Variation, expressed in percentage and estimated as the standard deviation of the parameter divided by the fixed effect and multiplied by 100. ***σ*** is vector of the residual error model parameters. Last column shows the relative standard errors (R.S.E.) of the estimates.

| Model | Parameter | Unit | Fixed effects | CV (%) | R.S.E. (%) |
| --- | --- | --- | --- | --- | --- |
| Reduced Gompertz | $\beta$ | day<sup>-1</sup> | 0.0771 | 95.8 | 4.21 |
| | $k$ | - | 9.85 | - | 0.935 |
| | $\sigma$ | - | [0,0.324] | - | [0,10.2] |
| Gompertz | $\alpha$ | day<sup>-1</sup> | 0.758 | 11.9 | 4.52 |
| | $\beta$ | day<sup>-1</sup> | 0.0768 | 128.4 | 5.69 |
| | $\sigma$ | - | [0,0.32] | - | [0,10.3] |
| Logistic | $\alpha$ | day<sup>-1</sup> | 0.405 | 12.1 | 2.39 |
| | $K$ | mm<sup>3</sup> | 1.19e+10 | 1.18e-9 | 8.78 |
| | $\sigma$ | - | [0,0.476] | - | [0,11.7] |
| Exponential | $\alpha$ | day<sup>-1</sup> | 0.077 | 39.98 | 9.58 |
| | $\sigma$ | - | [0,596] | - | [0,17.1] |

**Figure S1.**
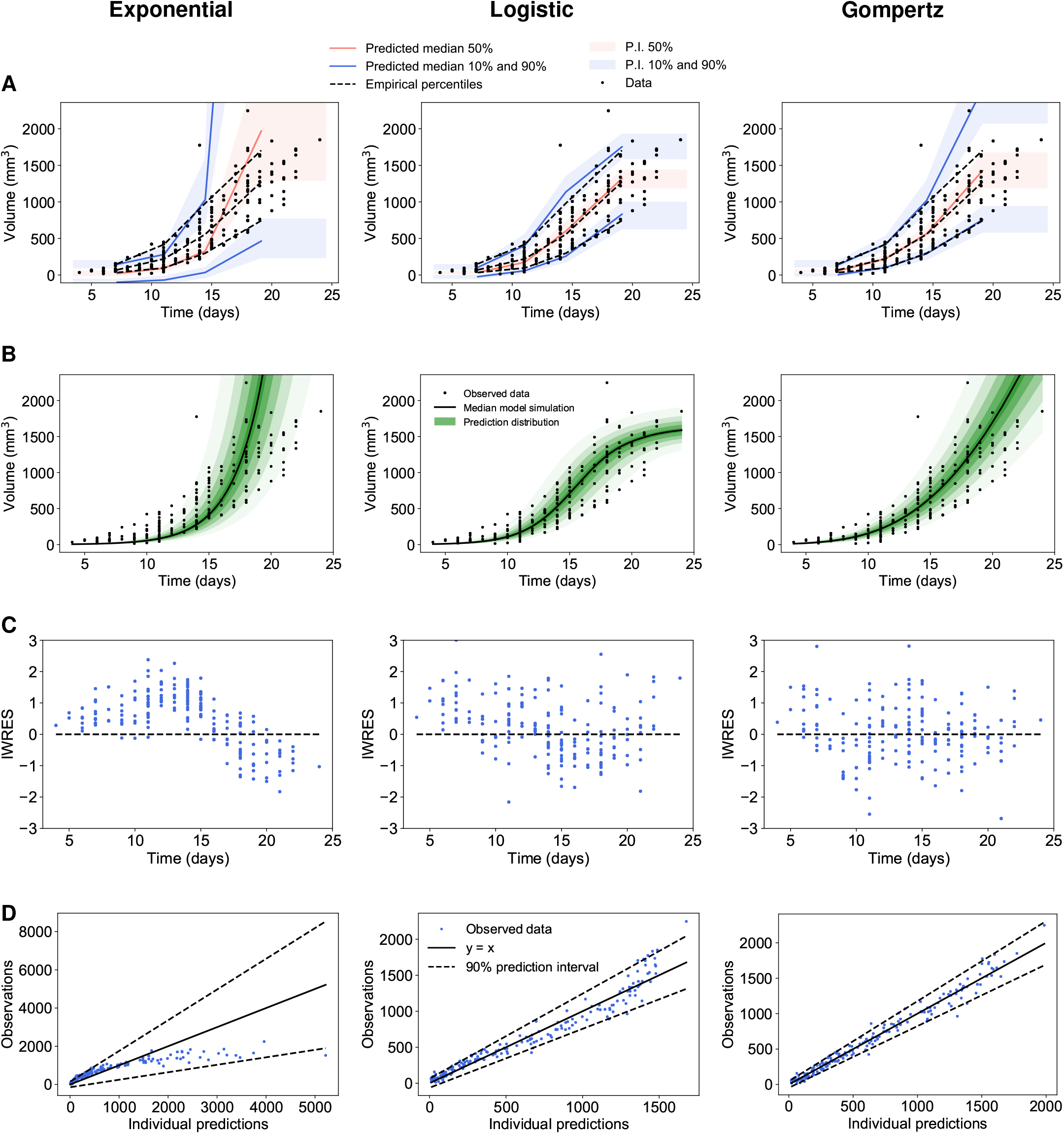
(lung). Diagnostic plots from population analysis. **Lung, volume.** Population analysis of experimental tumor growth kinetics. A) Visual predictive checks assess goodness-of-fit for both structural dynamics and inter-animal variability by reporting model-predicted percentiles (together with confidence prediction intervals (P.I) in comparison to empirical ones. B) Prediction distributions. C) Individual weighted residuals (IWRES) with respect to time. D) Observations vs predictions Left: exponential, Center: logistic, Right: Gompertz models.

**Figure S2.**
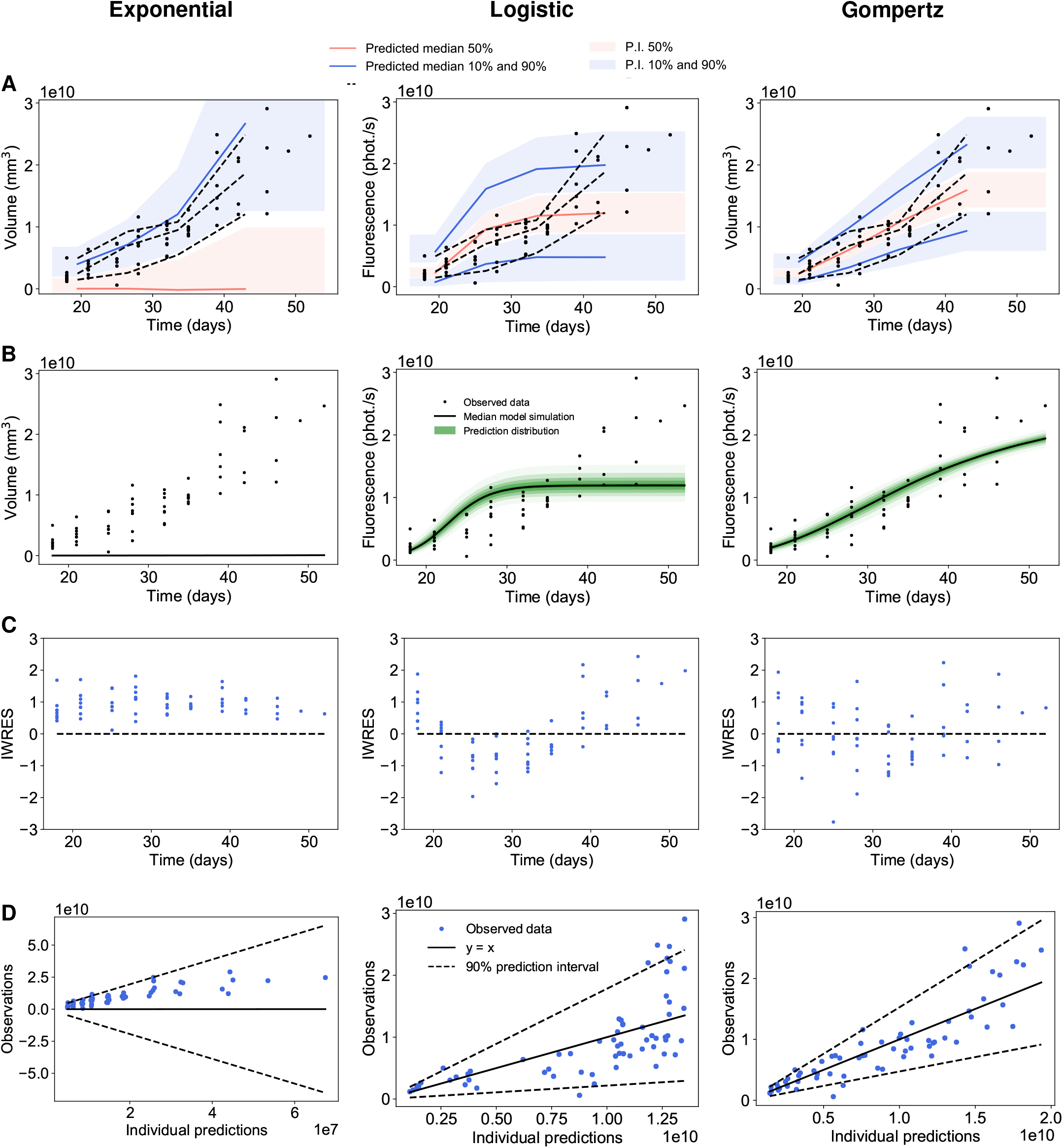
(breast-fluorescence). Diagnostic plots from population analysis. **Breast, fluorescence.** Population analysis of experimental tumor growth kinetics. A) Visual predictive checks assess goodness-of-fit for both structural dynamics and inter-animal variability by reporting model-predicted percentiles (together with confidence prediction intervals (P.I) in comparison to empirical ones. B) Prediction distributions. C) Individual weighted residuals (IWRES) with respect to time. D) Observations vs predictions Left: exponential, Center: logistic, Right: Gompertz models.

**Figure S3.**
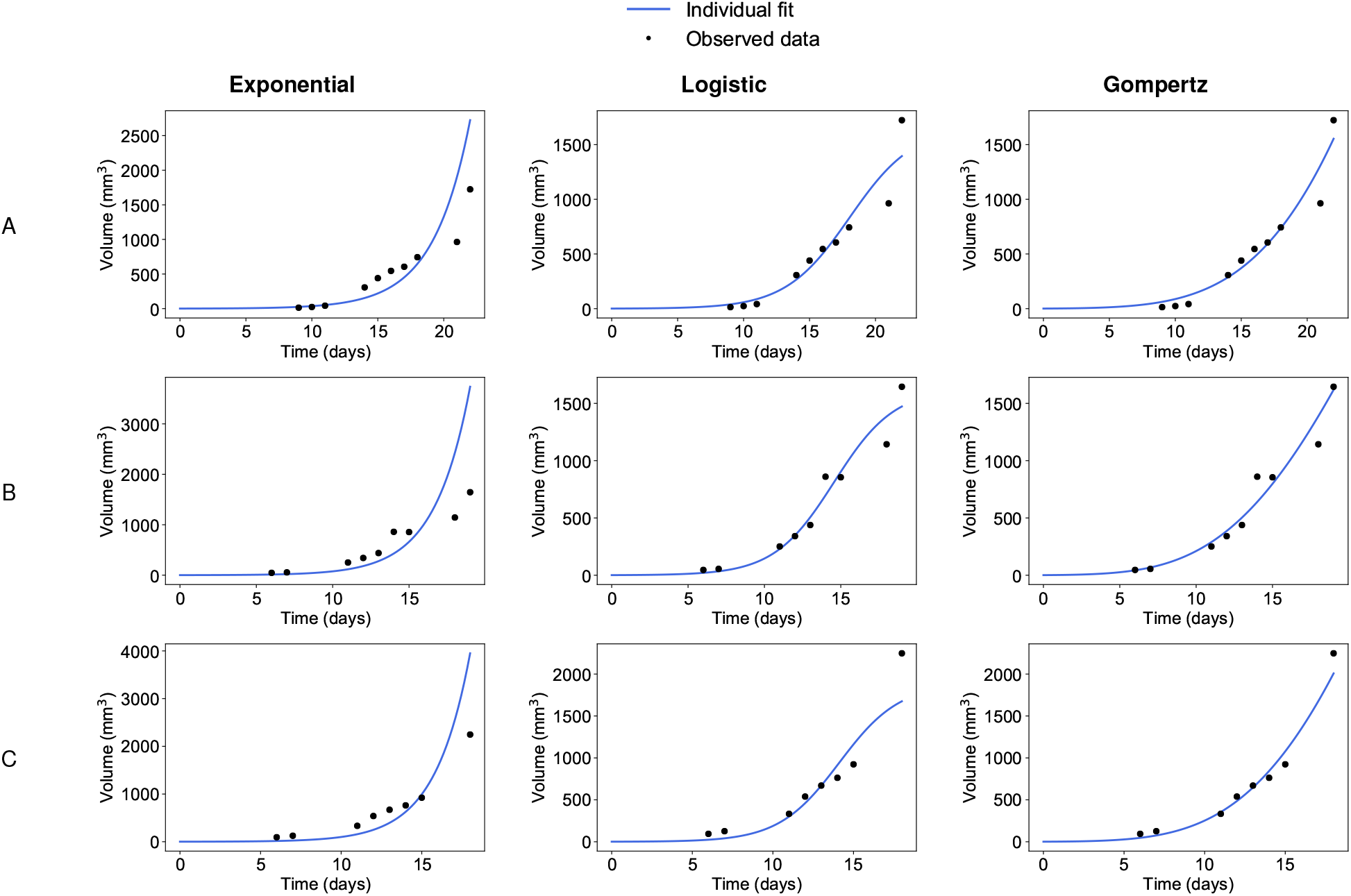
(lung). Individual fits from population analysis. **Lung, volume.** Three representative examples of individual fits computed with the population approach relative to the exponential (left), the logistic (center) and the Gompertz (right) models.

**Figure S4.**
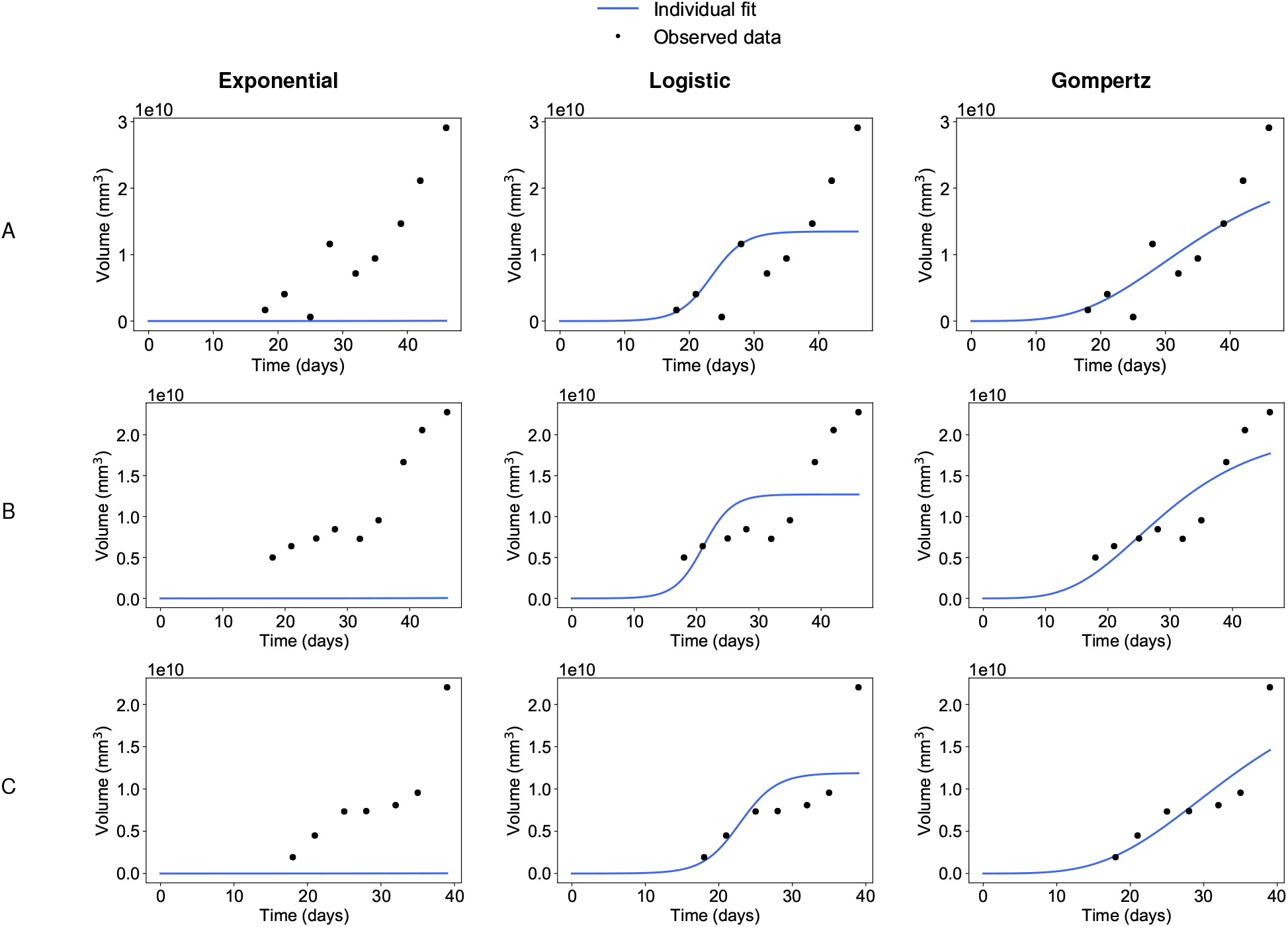
(breast-fluorescence). Individual fits from population analysis. **Breast, fluorescence.** Three representative examples of individual fits computed with the population approach relative to the exponential (left), the logistic (center) and the Gompertz (right) models.

### The reduced Gompertz model

**Figure S5.**
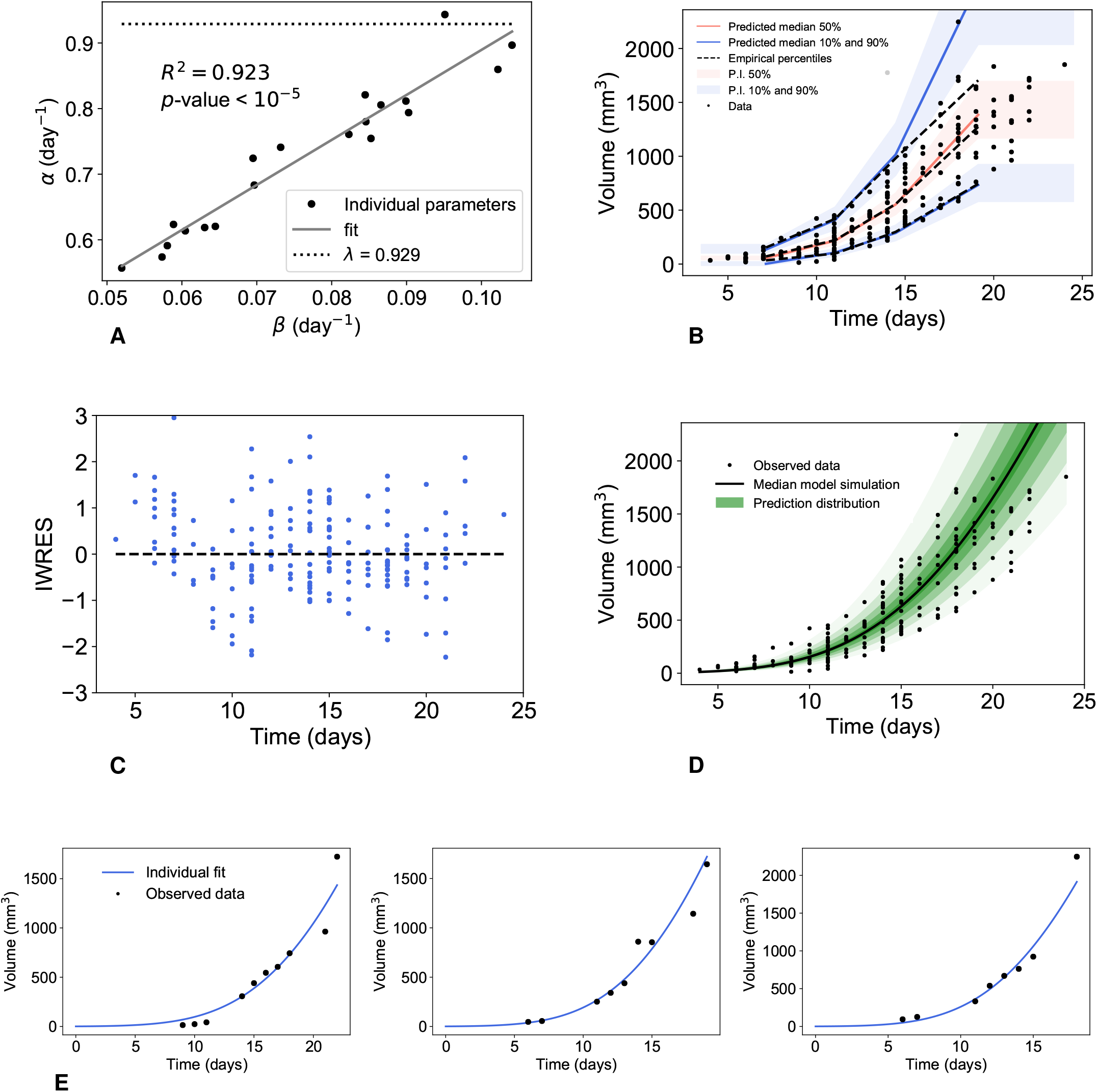
(lung). Correlation between the Gompertz parameters and diagnostic plots of the reduced Gompertz model with the population approach. **Lung, volume.** Correlation between the individual parameters of the Gompertz model (A) and results of the population analysis of the reduced Gompertz model: visual predictive check (B), examples of individual fits (C) and scatter plots of the residuals (D).

**Figure S6.**
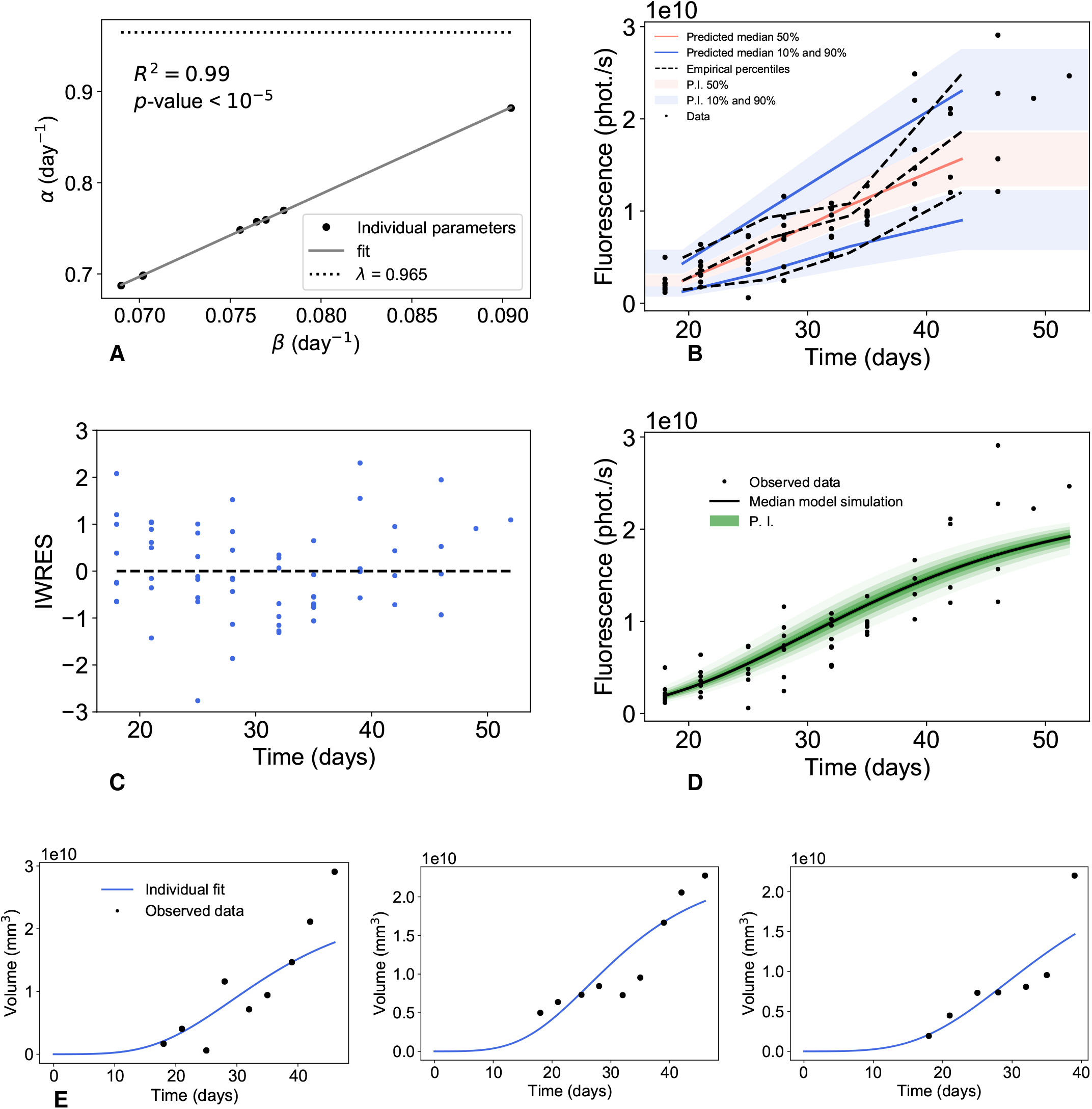
(breast-fluorescence). Correlation between the Gompertz parameters and diagnostic plots of the reduced Gompertz model with the population approach. **Breast, fluorescence.** Correlation between the individual parameters of the Gompertz model (A) and results of the population analysis of the reduced Gompertz model: visual predictive check (B), examples of individual fits (C) and scatter plots of the residuals (D).

### Prediction of the age of a tumor

**Figure S7.**
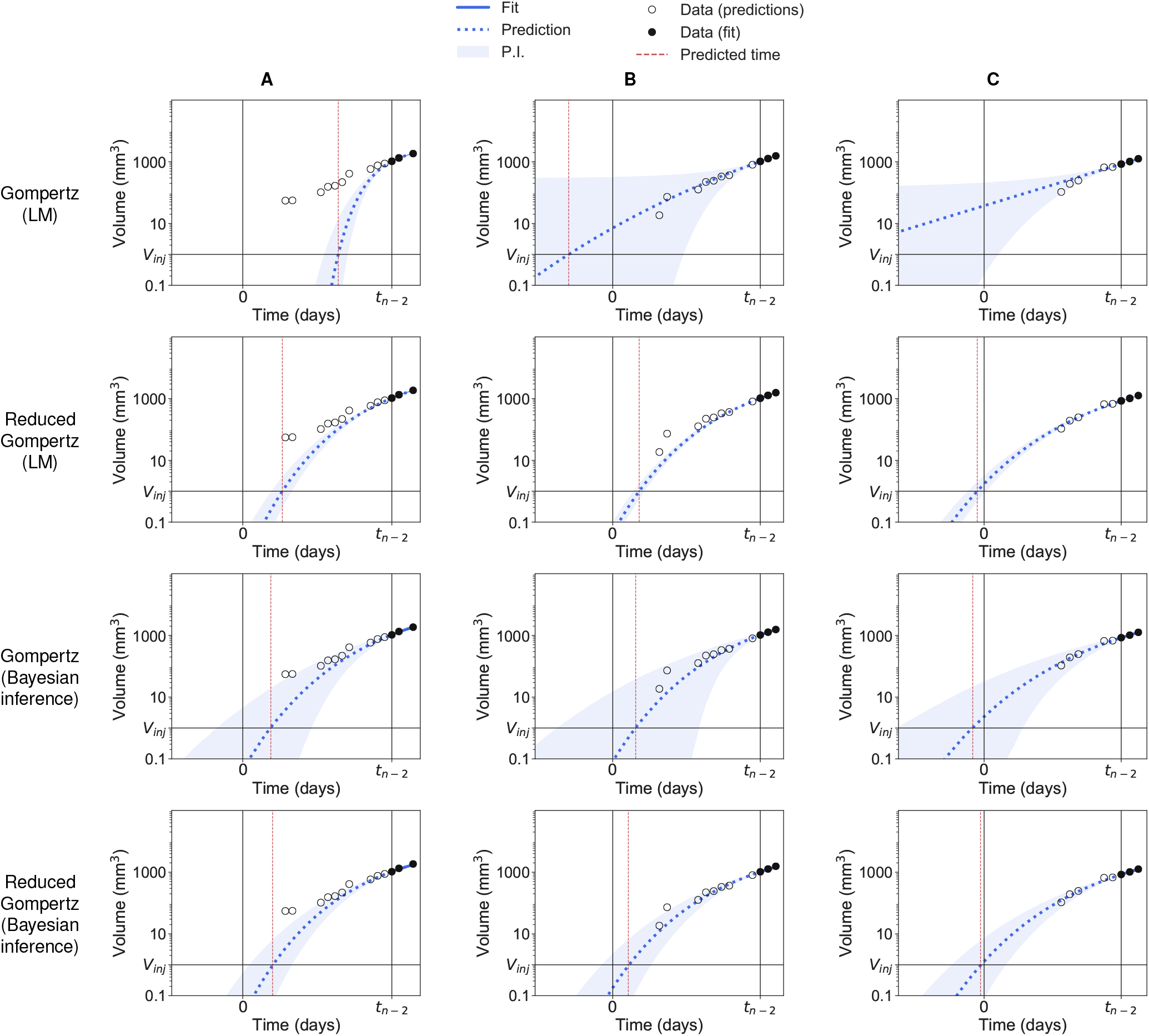
(lung). Backward predictions computed with likelihood maximization (LM) and with bayesian inference. **Lung, volume.** Three examples of backward predictions of individuals A, B and C computed with likelihood maximization (LM) and bayesian inference: Gompertz model with likelihood maximization (first row); reduced Gompertz with likelihood maximization (second row); Gompertz with bayesian inference (third row) and reduced Gompertz with bayesian inference (fourth row). Only the last three points are considered to estimate the parameters. The grey area is the 90% prediction interval (P.I) and the dotted blue line is the median of the posterior predictive distribution. The red line is the predicted initiation time and the black vertical line the actual initiation time.

**Figure S8.**
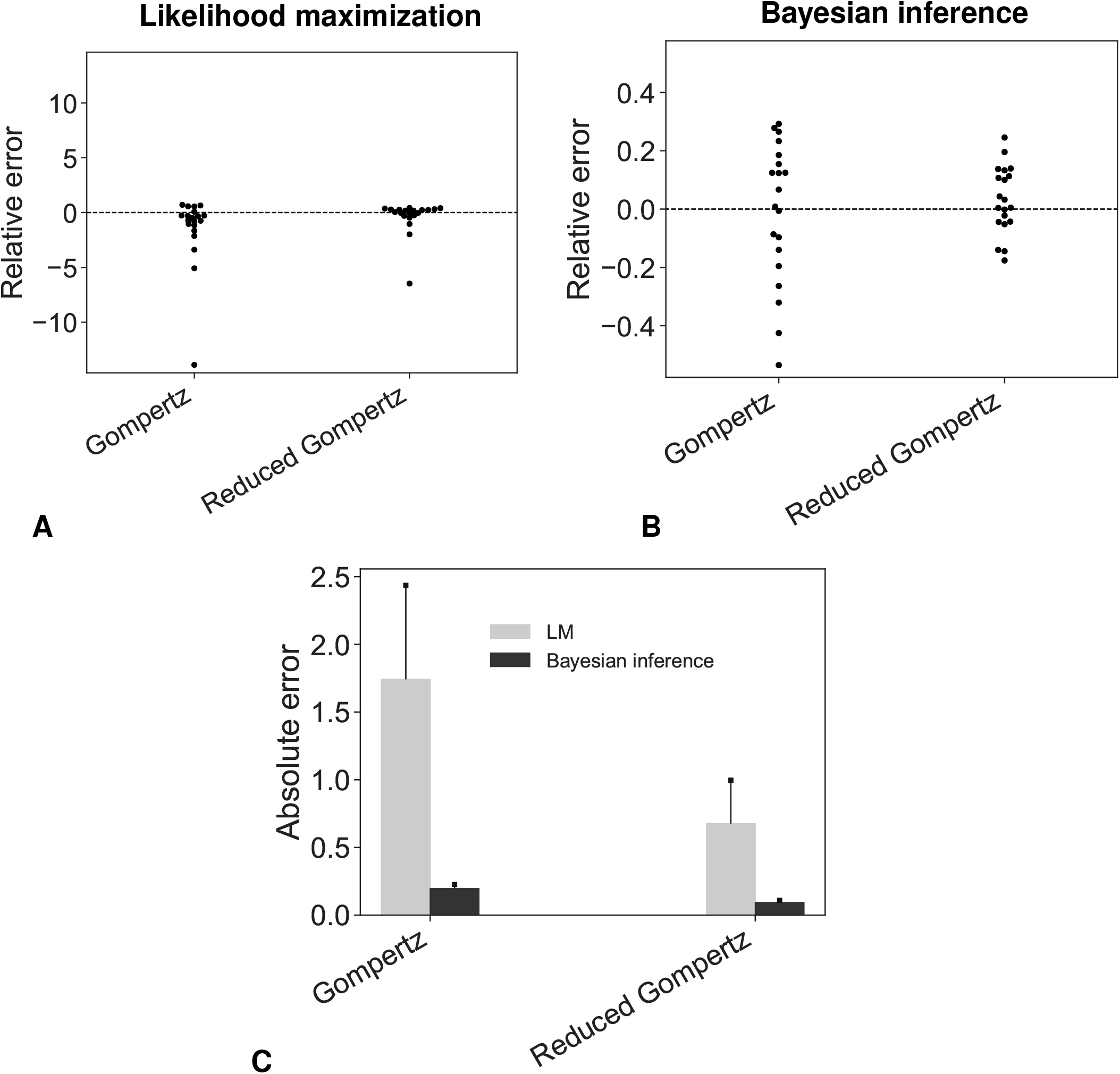
(lung). Error analysis of the predicted initiation time. **Lung, volume.** Accuracy of the prediction models. Swarmplots of relative errors obtained under likelihood maximization (A) or bayesian inference (B). (C) Absolute errors.

**Figure S9.**
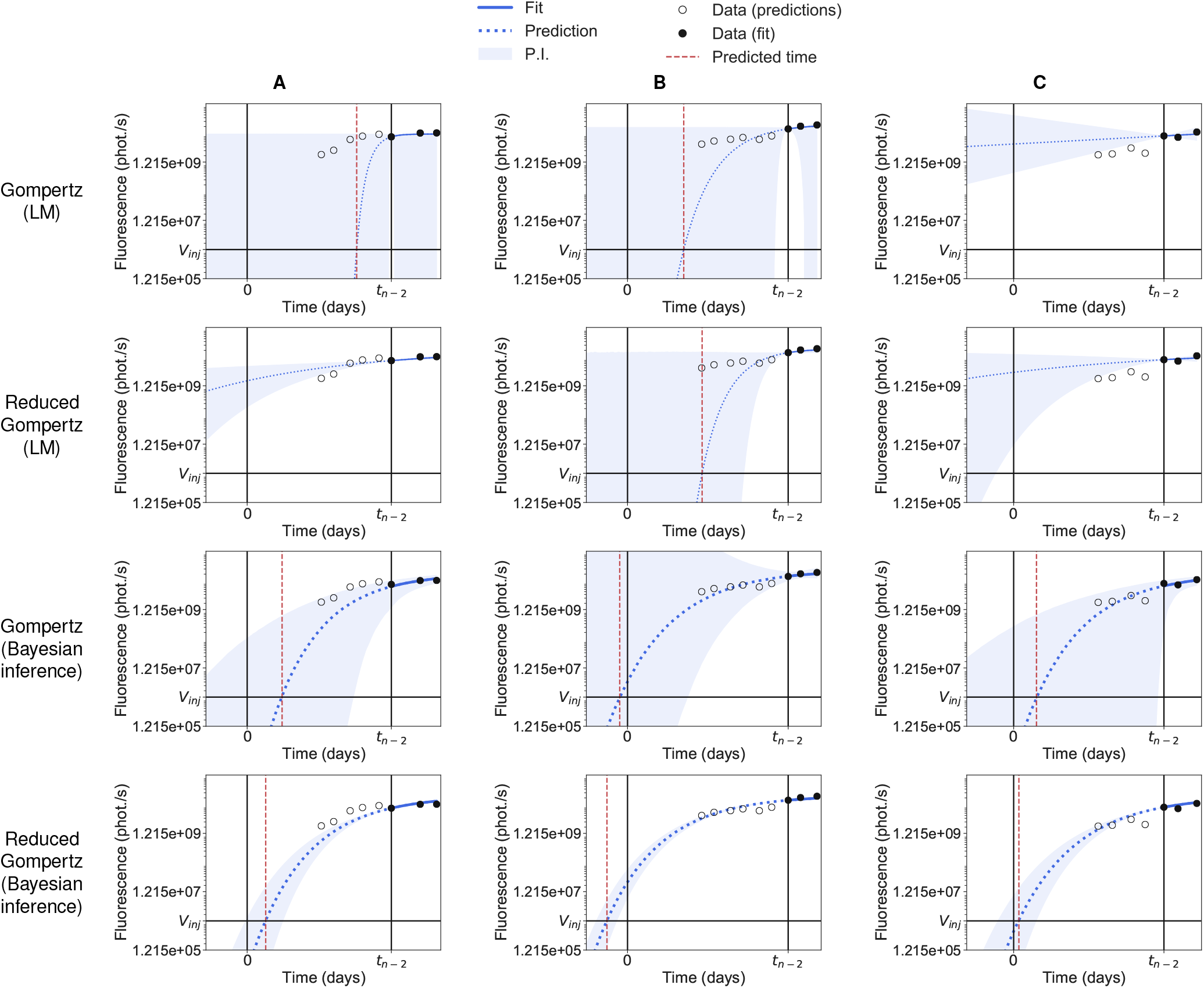
(breast-fluorescence). Backward predictions computed with likelihood maximization (LM) and with bayesian inference. **Breast, fluorescence.** Three examples of backward predictions of individuals A, B and C computed with likelihood maximization (LM) and bayesian inference: Gompertz model with likelihood maximization (first row); reduced Gompertz with likelihood maximization (second row); Gompertz with bayesian inference (third row) and reduced Gompertz with bayesian inference (fourth row). Only the last three points are considered to estimate the parameters. The grey area is the 90% prediction interval (P.I) and the dotted blue line is the median of the posterior predictive distribution. The red line is the predicted initiation time and the black vertical line the actual initiation time.

**Figure S10.**
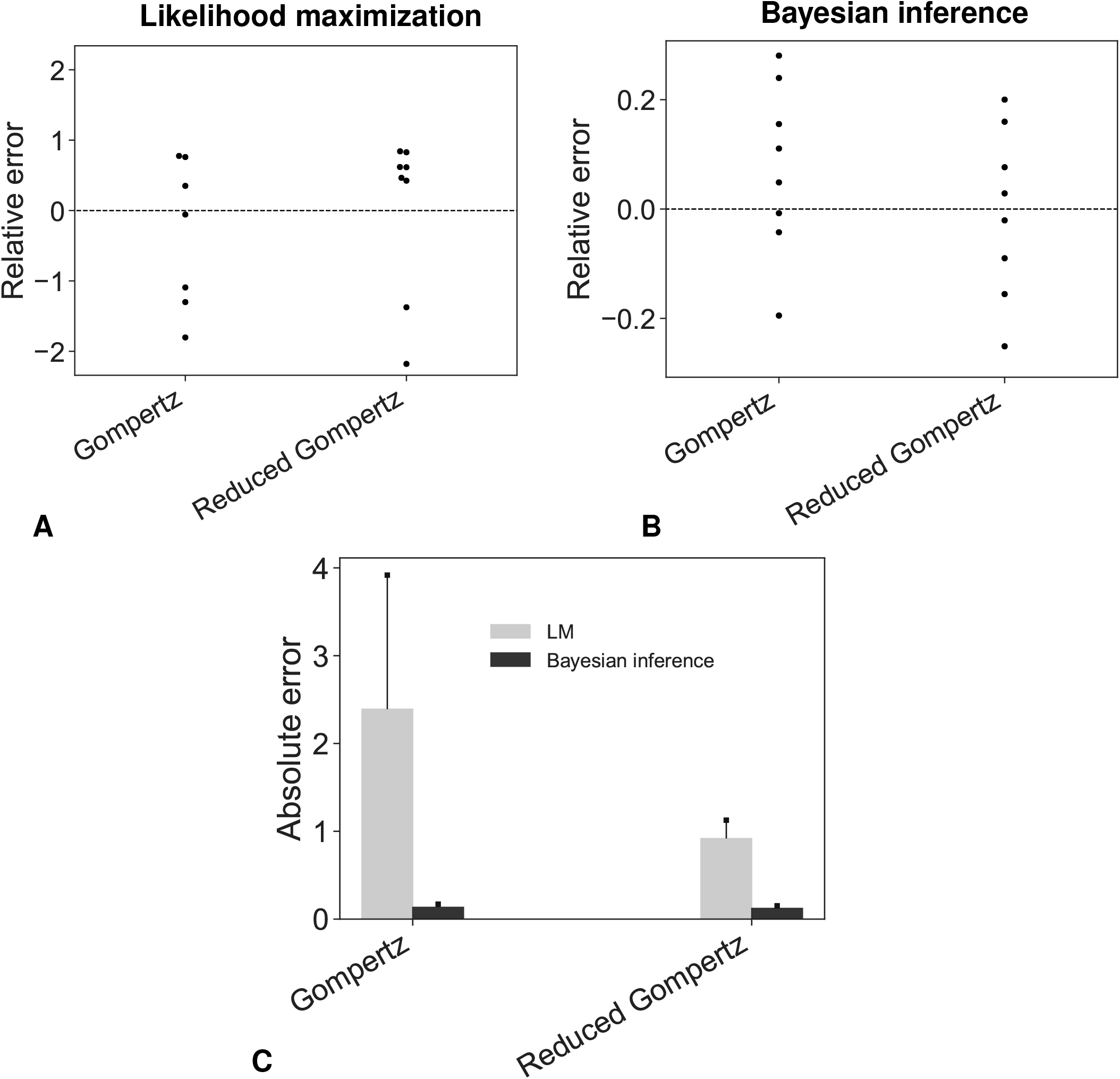
(breast-fluorescence). Error analysis of the predicted initiation time. **Breast, fluorescence.** Accuracy of the prediction models. Swarmplots of relative errors obtained under likelihood maximization (A) or bayesian inference (B). (C) Absolute errors.

